# Cohesin supercoils DNA during loop extrusion

**DOI:** 10.1101/2024.03.22.586228

**Authors:** Iain F. Davidson, Roman Barth, Sabrina Horn, Richard Janissen, Kota Nagasaka, Gordana Wutz, Roman R. Stocsits, Benedikt Bauer, Cees Dekker, Jan-Michael Peters

## Abstract

Cohesin extrudes genomic DNA into loops that promote chromatin assembly, gene regulation and recombination. Here we show that cohesin introduces negative supercoils into extruded DNA. Supercoiling requires engagement of cohesin’s ATPase heads, DNA clamping by these heads, and a DNA binding site on cohesin’s hinge, indicating that cohesin supercoils DNA when constraining it between the hinge and the clamp. Our results suggest that DNA extrusion stops once cohesin reaches its stall torque during supercoiling, and a cohesin mutant predicted to stall at lower torque forms shorter loops in cells. These results indicate that supercoiling is an integral part of the loop extrusion mechanism and that cohesin controls genome architecture not only by looping DNA but also by supercoiling it.

## Main Text

In eukaryotic interphase cells, the SMC (‘structural maintenance of chromosomes’) complex cohesin folds genomic DNA into loops and topologically associating domains (TADs; ref. (*1-4*)), which can regulate transcription (*5*), recombination (*6, 7*), sister chromatid separation (*8*) and replication (*9*). Cohesin extrudes DNA into loops (*10, 11*) through conformational changes that are controlled by ATP binding-hydrolysis cycles (*12*) (reviewed in (*13*)). These are catalyzed by cohesin’s SMC1 and SMC3 subunits, which contain 50 nm-long coiled-coils, dimerization ‘hinge’ domains and globular ATPase ‘heads’ (fig. S1A), which are related to those of ABC transporters (*14*). Upon ATP binding, cohesin’s heads engage and a subunit called NIPBL ‘clamps’ DNA on top of the engaged ATPase heads (ref. (*12, 15-17*); fig. S1B). These movements generate ∼15 pN force (*18*) and loop extrusion steps of ∼40 nm (100-200 bp; ref. (*19*)), indicating that DNA is reeled into the forming loop during head engagement.

In contrast, little is known about conformational changes of DNA during loop extrusion. Topoisomerase II binds and cleaves DNA at the base of cohesin loops (*20-23*), suggesting that DNA is supercoiled at these sites. The mitotic SMC complex condensin also co-localizes and interacts with topoisomerases (*24-30*) and can supercoil DNA *in vitro* (*31-33*). It has been proposed that this process occurs during loop extrusion (*31, 33*), but cohesin was found not to supercoil DNA (*34*), even though cohesin and condensin use similar loop extrusion mechanisms (*13*). It has therefore remained unclear whether DNA is generally supercoiled during loop extrusion, or whether topoisomerases accumulate at the base of SMC loops because supercoils that are generated by transcription are trapped there, as has been proposed (*22*).

### Cohesin and NIPBL supercoil DNA

We therefore tested whether cohesin can supercoil DNA in the presence of NIPBL, which is essential for loop extrusion but was not included in earlier experiments (*34*). For this purpose, we used an assay which measures changes in the linking number (Lk) of plasmids (*32*). Lk describes the number of times one strand passes over the other and is the sum of helical turns (or twist, Tw) and superhelical turns (or writhe, Wr; called plectonemes if multiple writhes accumulate).

For topologically constrained DNA molecules, Lk is invariant, i.e. any increase in Tw is compensated for by a decrease in Wr and *vice versa*. However, Lk can be altered by topoisomerases, which introduce DNA breaks, allow the DNA strands to rotate around each other, and reseal the breaks. If cohesin supercoils DNA and thus generates topological stress (changes in Tw or Wr) that are relaxed by a topoisomerase, the plasmid’s Lk will change. This ΔLk affects the shape and hydrodynamic radius of plasmids and can therefore be detected by a change in their electrophoretic mobility (Fig. 1A).

**Fig. 1.**
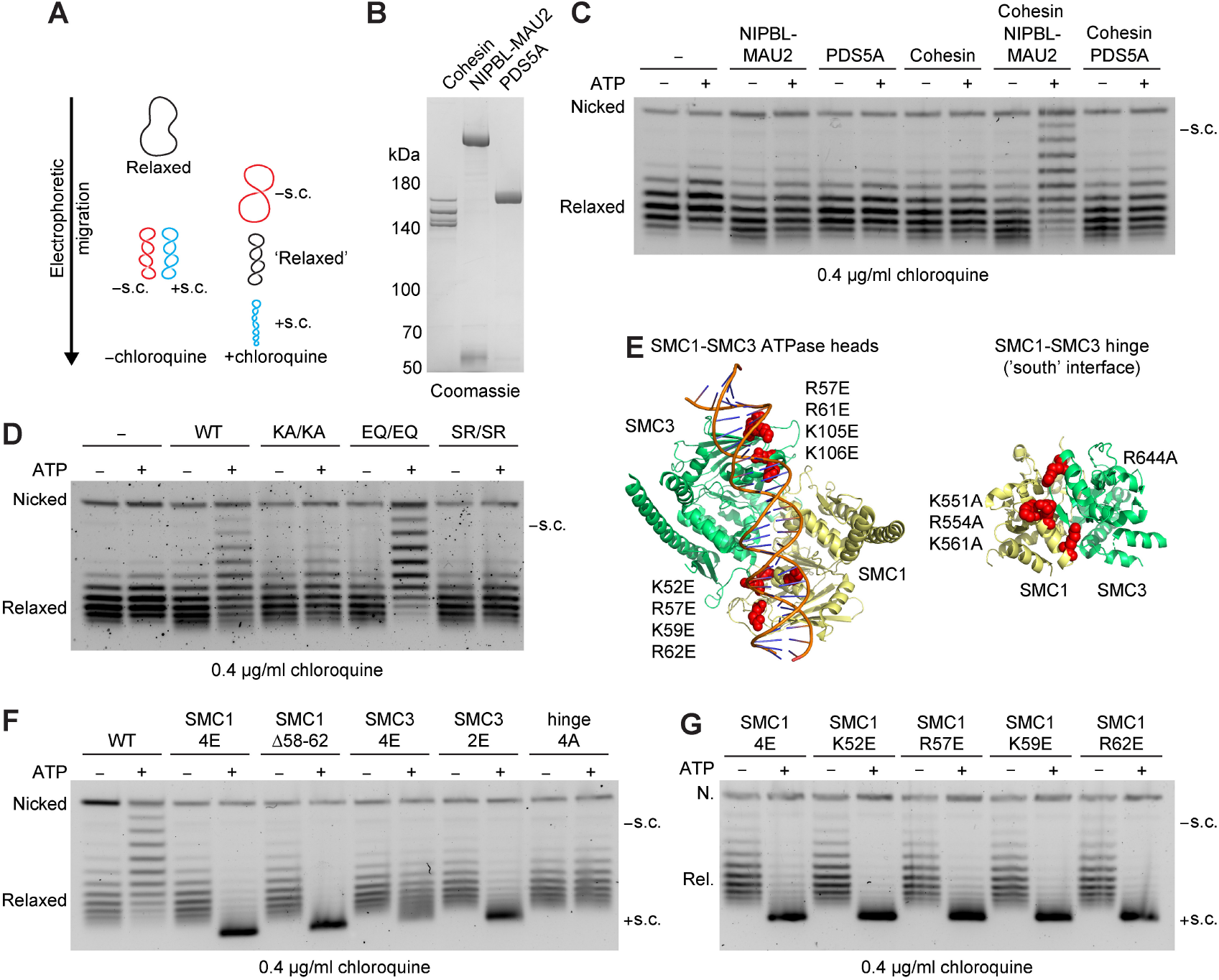
Human cohesin negatively supercoils DNA during clamping. **(A)** Chloroquine agarose gel electrophoresis allows discrimination between negatively and positively supercoiled plasmid DNA. **(B)** Coomassie staining of recombinant human cohesin, NIPBL-MAU2 and PDS5A after SDS-PAGE. **(C)** Human topoisomerase I DNA supercoiling assay. Negatively supercoiled plasmid DNA was relaxed by incubating with human topoisomerase I. Incubations were then supplemented as indicated with ATP, wild type cohesin at a 15:1 cohesin:plasmid ratio, NIPBL-MAU2 (30:1) or PDS5A (30:1). Reactions were terminated and purified DNA was separated by agarose gel electrophoresis in the presence of 0.4 μg/ml chloroquine. Representative image from 3 independent experiments. **(D)** As (C), except wild type, K38A/K38A, E1157Q/E1144Q or S1129R/S1116R cohesin was added at a 20:1 cohesin:plasmid ratio along with NIPBL-MAU2 (40:1). Representative image from 2 independent experiments. **(E)** Structures of the SMC1-SMC3 ATPase heads (upper panel; PDB: 6WG3) and the SMC1-SMC3 hinge (lower panel; PDB: 2WD5). DNA binding site mutations are indicated as spheres (*12*). **(F)** As (C), except wild type, SMC1^4E^ (K52E, R57E, K59E, R62E), SMC1^Δ58-62^, SMC3^4E^ (R57E, R61E, K105E, K106E), SMC3^2E^ (R57E, R61E) or hinge 4A (SMC1 K551A, R554A, K561A; SMC3 R644A) cohesin was added at a 15:1 molar ratio relative to plasmid DNA along with NIPBL-MAU2 (30:1). Representative image from 2 independent experiments. **(G)** As (C), except the indicated forms of cohesin were added at a 15:1 molar ratio relative to plasmid DNA along with NIPBL-MAU2 (30:1). Representative image from 2 independent experiments.

Supercoiling can occur in two directions. In relaxed DNA, the two strands twist around its helical axis in a right-handed manner once every ∼ 10.5 bp (ref. (*35*)). DNA molecules that are more or less twisted than this are ‘positively’ or ‘negatively’ supercoiled, respectively. The handedness of supercoiling can be detected by electrophoresis in the presence of the DNA intercalator chloroquine. Chloroquine induces positive supercoiling and thus causes positively supercoiled plasmids to migrate faster than relaxed DNA, and negatively supercoiled DNA to migrate slower (*36*) (Fig. 1A).

To analyze cohesin’s supercoiling activity, we used human topoisomerase I (fig. S1C, D), which relaxes negative and positive supercoils. In this assay, human tetrameric cohesin (containing SMC1, SMC3, SCC1 and STAG1; Fig. 1B and fig. S1E – I), did not alter the electrophoretic mobility of relaxed plasmids (Fig. 1C), as reported (*34*). However, when NIPBL-MAU2 (Fig. 1B) was added, more slowly migrating DNA bands appeared in an ATP-dependent manner (Fig. 1C, fig. S2), indicating that cohesin generates negatively supercoiled plasmids in the presence of NIPBL-MAU2. This was not the case when NIPBL-MAU2 was replaced by PDS5A (Fig.1B, C). Since cohesin’s loop extrusion activity also depends on ATP and NIPBL-MAU2 and is not supported by PDS5A (*10, 11*), these results suggest that cohesin supercoils DNA during loop extrusion. Unless otherwise stated, all subsequent supercoiling experiments were therefore performed in the presence of NIPBL-MAU2.

### Cohesin supercoils DNA during clamping

To understand at which step of its ATP binding-hydrolysis cycle cohesin supercoils DNA, we analyzed the effect of mutations in the Walker A motifs (K38A/K38A, hereafter KA/KA), the signature motifs (S1129R/S1116R; SR/SR), or the Walker B motifs (E1157Q/E1144Q; EQ/EQ) of SMC1 and SMC3. These mutations block ATP binding, head dimerization and ATP hydrolysis, respectively (*14, 37*). We found that supercoiling was inhibited by KA/KA and SR/SR but not by the EQ/EQ mutations (Fig. 1D, fig. S2), indicating that supercoiling depends on ATP-dependent head engagement but not on subsequent ATP hydrolysis. Since cryogenic electron microscopy (cryo-EM; ref. (*15-17*)) and single-molecule Förster resonance energy transfer experiments (smFRET; ref. (*12*)) have shown that EQ/EQ mutations stabilize the head-engaged state of cohesin in which DNA is clamped onto the ATPase heads by NIPBL, these results suggest that cohesin induces supercoiling during DNA clamping.

To test this hypothesis, we analyzed cohesin in which the DNA binding sites on the ATPase heads of SMC1 (SMC1^4E^) or SMC3 (SMC3^4E^) were mutated (ref. (*12*); Fig. 1E) and which are therefore predicted to be defective in DNA clamping. Indeed, no slowly migrating DNA bands were detected in the presence of these complexes, suggesting that cohesin mutants that are defective in DNA clamping are also defective in generating negatively supercoiled DNA (Fig. 1F).

However, these experiments revealed an unexpected phenomenon. Supercoiling reactions containing cohesin-SMC1^4E^ were not only defective in generating slowly migrating DNA, but instead greatly increased the electrophoretic mobility of plasmids in an ATP and topoisomerase I dependent manner, indicating that they became positively supercoiled (Fig. 1F and fig. S2). A five amino acid deletion in SMC1 that includes two of the residues mutated in SMC1^4E^ (SMC1^Δ58-62^) had a similar effect (Fig. 1F). Remarkably, charge reversal of K52, R57, K59 and R62 alone also caused positive supercoiling (Fig. 1G). These residues are located in a loop that contains the ‘arginine finger’ R57, which is required for the DNA-dependent ATPase activity of *B. subtilis* SMC (R57 in human SMC1 corresponds to R59 in *B. subtilis* SMC; ref. (*38*)).

Fast migrating DNA was also observed after incubation with cohesin-SMC3^2E^ and to a lesser extent with cohesin-SMC3^4E^ (Fig. 1F). The overall reduction of supercoiling in the presence of cohesin-SMC3^4E^ is presumably due to the fact that this mutant binds DNA even less well than cohesin-SMC1^4E^ and cohesin-SMC3^2E^, as indicated by their DNA dependent ATPase activities (*12*). In contrast, mutation of a DNA binding site on cohesin’s hinge (hinge^4A^) completely abolished cohesin’s ability to generate either negatively or positively supercoiled plasmids (Fig. 1F), indicating that this DNA binding site is also required for supercoiling.

These results suggest that weakening the DNA binding sites of the clamp results in a switch from negative to positive supercoiling, whereas reducing DNA binding more strongly either in the clamp or at the hinge prevents supercoiling. These results also support the notion that cohesin supercoils DNA during loop extrusion, since all mutants defective in negative supercoiling are also reduced in their ability to extrude DNA (*12*).

### Visualization of cohesin-mediated DNA supercoiling using HS-AFM

To confirm that plasmids incubated with cohesin and topoisomerase I become supercoiled, we re-isolated these plasmids, visualized them by high-speed AFM (HS-AFM) and counted the number of DNA crossings per plasmid (Fig. 2, fig. S3 and movies S1 and S2). Plasmids that had previously been incubated with cohesin displayed a higher number of DNA crossings, and cohesin^EQ/EQ^ and cohesin-SMC1^4E^ induced more DNA crossings than wild type cohesin (Fig. 2C–E and fig.S3). These results support the interpretation that wild type cohesin, cohesin^EQ/EQ^, and cohesin-SMC1^4E^ all induce supercoiling (where we note that negatively and positively supercoiled plasmids cannot be distinguished in these HS-AFM data).

**Fig. 2.**
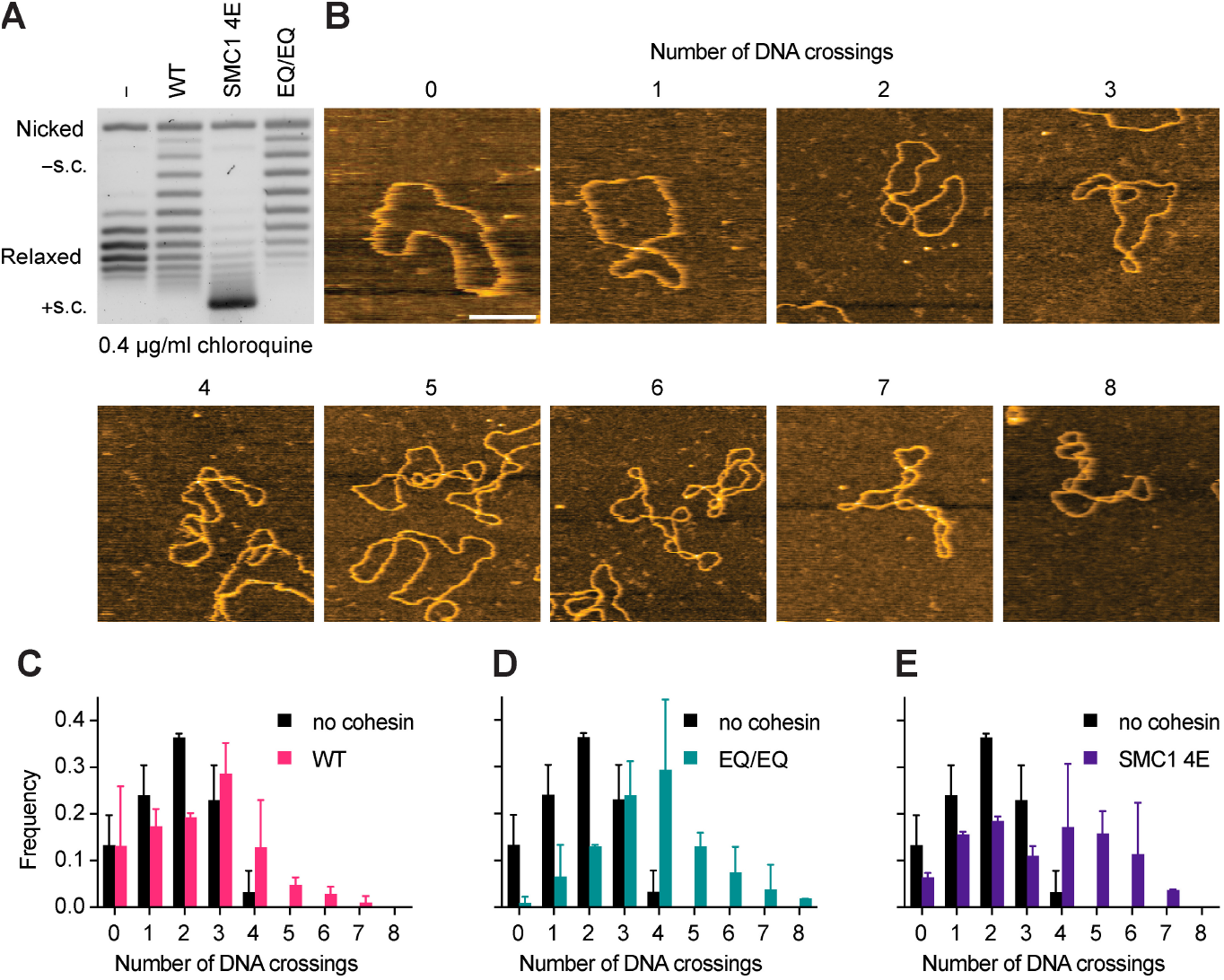
Visualization of supercoiled DNA by HS-AFM. **(A)** Human topoisomerase I DNA supercoiling assay in the presence or absence of the indicated proteins and ATP. Cohesin was added at a 15:1 molar ratio relative to plasmid DNA. NIPBL-MAU2 was added to all reactions at a 30:1 molar ratio relative to plasmid DNA. **(B – E)** Purified DNA from (A) was visualized using high-speed atomic force microscopy (HS-AFM). (B) Images of plasmid DNA with the indicated number of DNA crossings. The images with 0 – 6 DNA crossings were recorded using the EQ/EQ DNA sample shown in (A). The image with 7 DNA crossings was recorded using the SMC1 4E DNA sample shown in (A). (C – E) Frequency of the number of DNA crossings observed per DNA molecule by HS-AFM. Data are mean ± s.d. from two independent experiments. N = 102, 104, 108, 108 plasmids analyzed per no cohesin, wild type cohesin, cohesin-SMC1^4E^ and cohesin^EQ/EQ^ condition, respectively.

### Analysis of plasmid supercoiling handedness

We next tested whether the plasmids generated in the presence of cohesin-SMC1^4E^ were indeed positively supercoiled, as we assumed in our interpretations above. Notably, highly negatively supercoiled plasmids can also have an increased electrophoretic mobility if the number of negative supercoils exceeds the number of positive supercoils induced by chloroquine (fig. S1D). We therefore re-isolated plasmids supercoiled in the presence of topoisomerase I and cohesin-SMC1^4E^ or cohesin^EQ/EQ^ and treated them with topoisomerases, which preferentially relax positive supercoils (Topo IV), introduce negative supercoils (gyrase), or relax negative supercoils (TopA). If the DNA supercoils generated in the presence of cohesin-SMC1^4E^ and cohesin^EQ/EQ^ are indeed of opposite handedness, these plasmids should show differential sensitivity to these enzymes. Indeed, all experiments performed with Topo IV, DNA gyrase and TopA confirmed that plasmids become negatively supercoiled in the presence of cohesin^EQ/EQ^ but positively supercoiled in the presence of cohesin-SMC1^4E^ (fig. S4, see Supplementary Text).

Our observation that mutation of cohesin’s DNA clamping sites results in a switch from negative to positive supercoiling is reminiscent of the recent observation that plasmids become negatively supercoiled at low condensin-to-DNA ratios but positively supercoiled at high ratios (*33*). Indeed, we observed a similar switch when cohesin-to-plasmid ratios were increased (Fig. 3A).

**Fig. 3.**
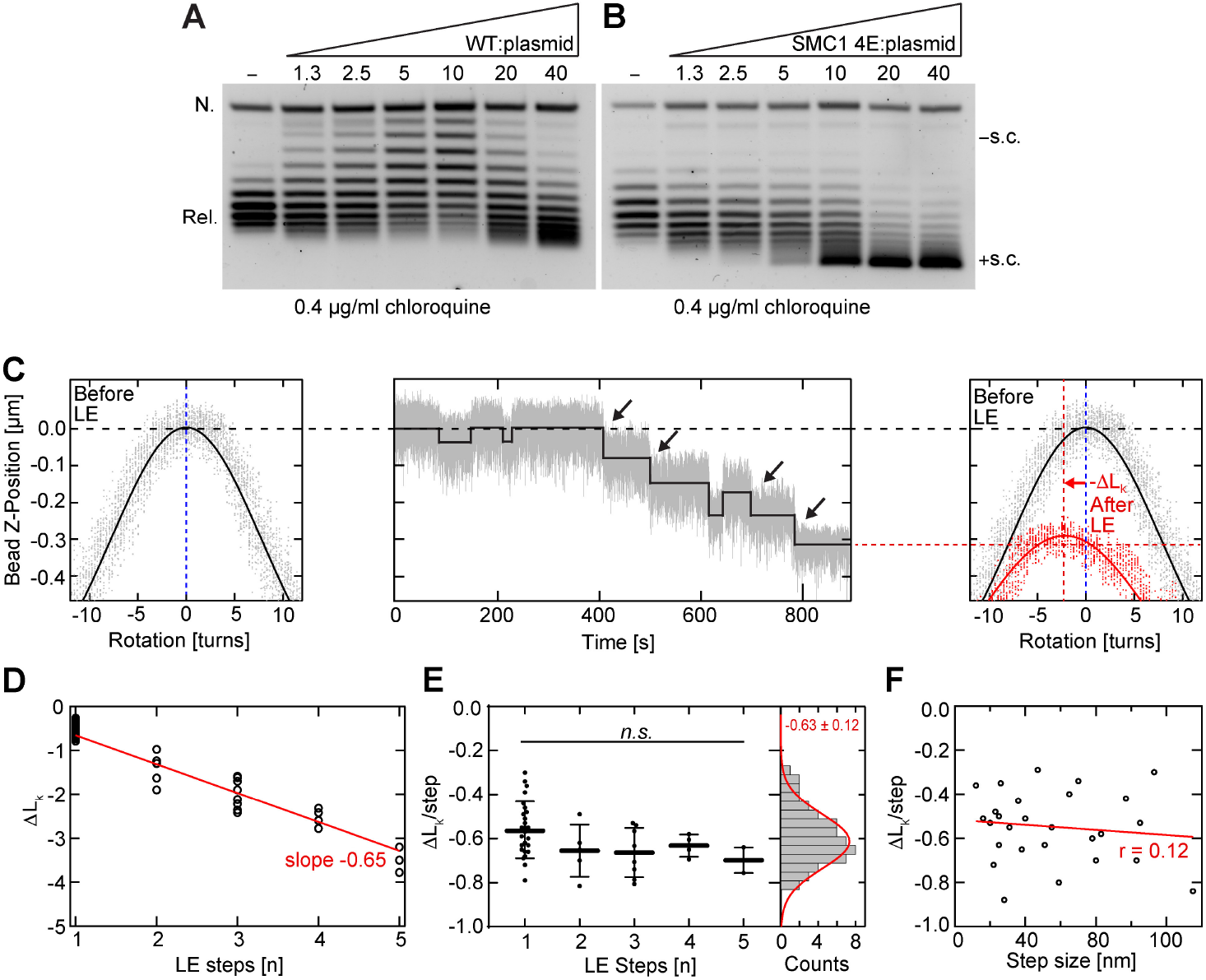
Individual loop extrusion steps by cohesin-SMC1^4E^ introduce negative twist into the extruded DNA loop. **(A – B)** Human topoisomerase I DNA supercoiling assay in the presence of ATP and (A) wild type cohesin or (B) cohesin-SMC1^4E^ at the indicated molar ratios relative to plasmid DNA. NIPBL-MAU2 was added to all reactions at an 80:1 molar ratio relative to plasmid DNA. Representative images from 2 independent experiments. **(C)** (Left) DNA end-to-end extension as a function of magnet rotation for a torsionally constrained 3.6 kbp DNA molecule. The DNA extension at constant 0.3 pN force is maximal when no external rotations are applied. Upon applying positive/negative rotations, the over/underwinding of the DNA changes the linking number Lk and leads to the formation of supercoils, which are at this force symmetric for both positive and negative coiling. The solid black line depicts a Gaussian fit to the rotation curve data. (Center) Representative trajectory of cohesin-SMC1^4E^ (20 pM cohesin-SMC1^4E^, 50 pM NIPBL-MAU2) showing step-wise DNA loop extrusion in the presence of 1 mM ATP at 0.3 pN. The black line depicts fit from the step-finding algorithm. (Right) DNA extension as a function of magnet rotation similar to (Left), conducted directly after the DNA loop extrusion experiment. The rotation curve after loop extrusion (red; solid line depicts Gaussian fit to the data) shows that the maximum DNA extension was shifted to negative magnet rotation compared to the initial rotation curve (black), caused by the uncoiling of the positive, complementary supercoils (+ΔLk) formed in the DNA molecule outside the DNA loop. The degree of negative supercoils (-ΔLk) generated by cohesin-SMC1^4E^ residing in the loop (red arrow) is equal to the degree of positive supercoils (+ΔLk) generated outside the loop. **(D)** Change in DNA linking number, ΔLk, over the number of steps for cohesin-SMC1^4E^ in the presence of ATP (total N= 54). The red line represents a linear fit without offset of the change in linking number over the number of LE steps. **(E)** The same data as in (D) but divided by the number of LE steps. The histogram on the right represents all data points, fitted by a Gaussian with -0.63 ± 0.12 turns (mean ± SD). Statistical significance was assessed using ANOVA with a significance level α = 0.05 (95% confidence interval; n.s. = p >0.05). **(F)** Change in rotation per step is plotted against the step size for traces with only a single step (N = 24). Linear fit and Pearson’s correlation coefficient are shown in red.

Because positively supercoiled plasmids are generated in the presence of high wild type cohesin-to-DNA ratios and by cohesin-SMC1^4E^ at any ratio, we wondered whether cohesin-SMC1^4E^ is hyperactive in supercoiling, i.e. whether it mimics high wild-type cohesin-to-DNA ratios. If so, one might observe negative supercoiling when cohesin-SMC1^4E^ is incubated at low cohesin-to-plasmid ratios. However, negatively supercoiled plasmids could neither be detected in dose-response (Fig. 3B) nor in time course (fig. S5A) analyses of reactions containing cohesin-SMC1^4E^, indicating that this mutant is not a hyperactive form of cohesin.

### Cohesin-SMC1^4E^ generates a negative twist of -0.6 in each loop extrusion step

The switch in supercoiling handedness observed with DNA clamping mutants and at high cohesin-to-plasmid ratios could represent a change in the handedness of cohesin’s supercoiling activity or could result from an indirect effect. To distinguish between these possibilities, we analyzed cohesin-SMC1^4E^ in a magnetic tweezers assay in which the degree of DNA twisting that occurs during individual loop extrusion steps can be measured (see the accompanying manuscript by Janissen et al, 2024). Cohesin-SMC1^4E^ was able to induce step-like DNA shortening events in this assay (Fig. 3C), suggesting that cohesin-SMC1^4E^ is able to carry out loop extrusion steps (Fig. 3C), even though this mutant displayed only little extrusion activity in fluorescence microscopy assays (3 % of wild type; ref. (*12*)).

Importantly, in this magnetic tweezers assay, cohesin-SMC1^4E^ generated a *negative* twist of - 0.63 ± 0.12 (mean ± SD) in each loop extrusion step (Fig. 3C–F), a value that has the same handedness and degree of twist as observed in the presence of wild type cohesin (Janissen et al., 2024). This suggests that the switch in supercoiling handedness does not represent a change in handedness with which cohesin-SMC1^4E^ supercoils DNA, but instead is caused by an indirect effect in the plasmid supercoiling assay.

### Simulations predict a switch in supercoiling handedness

To test whether this effect could be related to the fact that Lk in the plasmid supercoiling assay is altered by topoisomerase I, and not by cohesin, we used Monte Carlo simulations to model plasmid supercoiling *in silico* based on a set of simple parameters (table S1). We reasoned that loop extrusion causes differential changes in the supercoiling density in the extruded and the non-extruded segments of the plasmid (Fig. 4A). Since every loop extrusion step introduces the same degree of negative twist (Janissen et al., 2024) and reels a similar amount of DNA into the loop (*19*), the ratio of supercoils per DNA unit length, i.e. the supercoiling density, will initially remain constant in the growing DNA loop. In contrast, in the non-looped segment the supercoiling density will increase with every loop extrusion step since each of these will shorten the segment but simultaneously introduce compensatory positive twist.

**Fig. 4.**
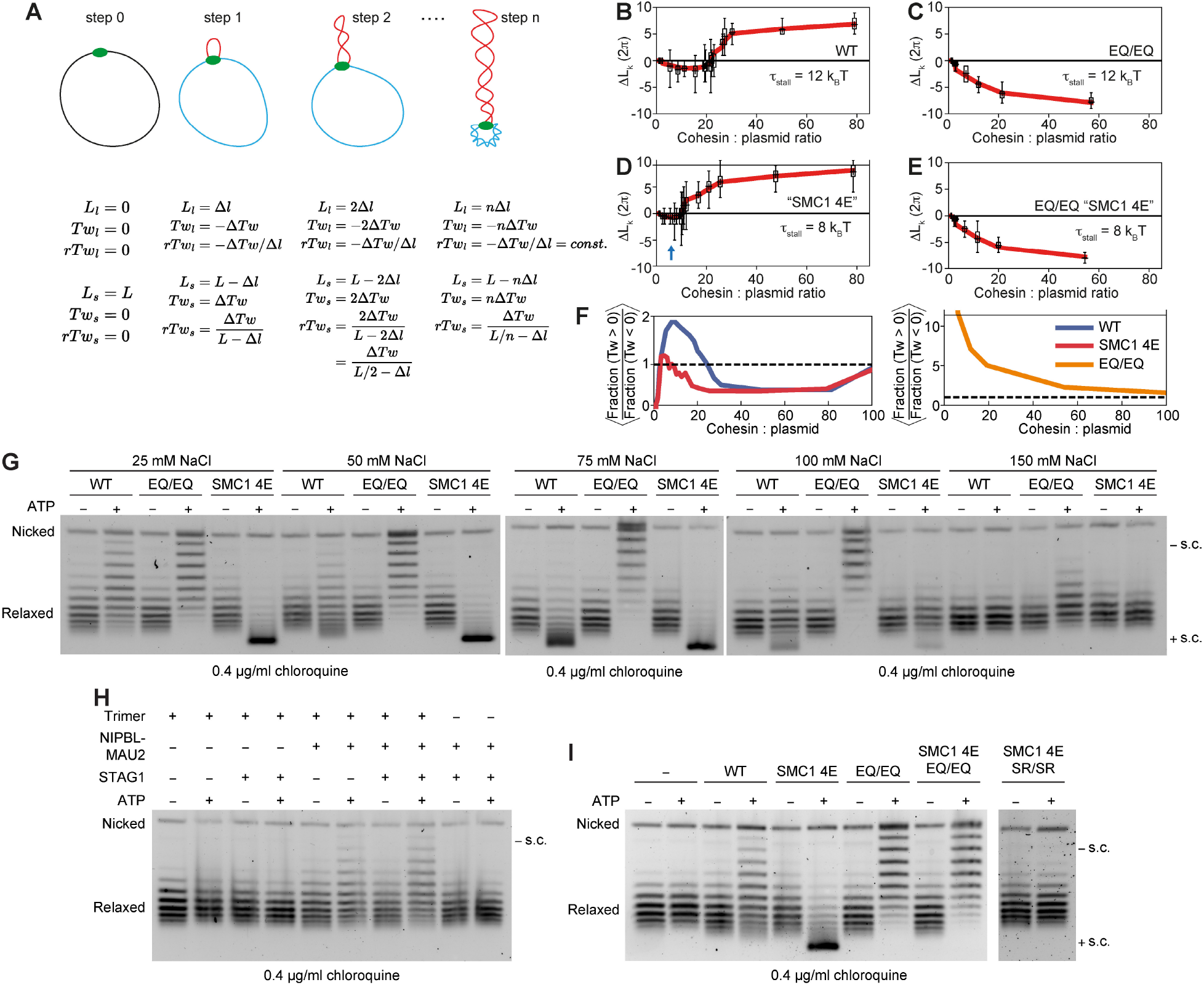
Loop extrusion-induced torque simulations can recapitulate the transition from negative to positive supercoiling chirality at high cohesin:plasmid DNA ratios. **(A)** Illustration of the twist (*Tw*), segment length (*L*), and relative twist (*rTw*) in the loop *l* and the plasmid segment *s* during the first n steps of loop extrusion by a single cohesin complex. The relative twist within the plasmid segment *s* increases with n while the relative twist (and thus supercoiling density and torque) within the loop remains approximately constant. **(B)** Simulation of the number and chirality of linking numbers removed (ΔLk) by human topoisomerase I at the indicated wild type cohesin:plasmid ratios. **(C)** As (B) but cohesin^EQ/EQ^ was simulated by allowing only one step per cohesin. **(D)** As (B) except the torque at which loop extrusion stalls (*t*_stall_) was set at 8 kT. **(E)** as (B) except cohesin-SMC1^4E-EQ/EQ^ was simulated by setting *t*_stall_ at 8 kT and allowing only one step per cohesin. Bar plots represent the mean value (+ sign), whiskers denote the span of the data. The red line is a guide to the eye and was constructed by applying a Savitzky-Golay filter of window length 5 and order 1. Data are from 50 independent simulations. **(F)** At every time point, the fraction of the plasmid with a positive twist (Fraction (Tw > 0)) is divided by the fraction of the plasmid with negative twist (Fraction (Tw < 0)). The quotient is then averaged over time and subsequently over 200 independent simulations and plotted at the indicated cohesin:plasmid ratios. For wild type cohesin, the majority of the plasmid is positively supercoiled for cohesin:plasmid ratios up to ∼30, while cohesin-SMC1^4E^ rarely produces a predominantly positively supercoiled plasmid. In contrast, cohesin^EQ/EQ^ can only make one step which yields that the majority of the plasmid is unlooped, carrying the positive compensatory twist from the single steps of the individual cohesin^EQ/EQ^ complexes. **(G)** Human topoisomerase I DNA supercoiling assay at different NaCl concentrations in the presence or absence of the indicated proteins and ATP. Cohesin was added at a 15:1 molar ratio relative to plasmid DNA. NIPBL-MAU2 was added to all reactions at a 30:1 molar ratio relative to plasmid DNA. Representative images from 2 independent experiments. **(H)** As (G), except trimeric cohesin was added at a 15:1 molar ratio relative to plasmid DNA in the presence or absence of NIPBL-MAU2 (30:1) or STAG1 (45:1) in a buffer containing 25 mM NaCl. Representative image from 4 independent experiments. **(I)** As (G) except the indicated forms of cohesin were added at a 20:1 cohesin:plasmid ratio in a buffer containing 25 mM NaCl. NIPBL-MAU2 was added to all reactions at a 40:1 molar ratio relative to plasmid DNA. Representative images from 2 independent experiments.

In this scenario, each loop extrusion step transfers a certain degree of positive supercoiling from the non-looped segment into the growing loop. Furthermore, cohesin could initiate extrusion in loops previously formed by other cohesin complexes, potentially resulting in complex patterns of non-extruded, extruded and re-extruded plasmid parts as well as Z-loops (*39*) with different length distributions and supercoiling states, in which the numbers, densities and handedness of supercoils could vary. Our simulations allowed these complex processes to occur.

Furthermore, we assumed that topoisomerase I binds to all parts of the plasmid equally well. In plasmids containing identical numbers of cohesin-induced negative supercoils and compensatory positive supercoils topoisomerase I would therefore preferentially bind to and resolve negative supercoils if the negative supercoils were distributed over a larger fraction of the plasmid than the positive supercoils. The opposite would be true if positive supercoils were distributed over a larger fraction.

Remarkably, we found that these simulations predicted a switch from negative to positive supercoiling exactly under the conditions we had experimentally observed (Fig. 4B–E and movies S3 and S4). For wild type cohesin, negative supercoiling was observed at low cohesin-to-plasmid ratios but positive supercoiling at high ratios (Fig. 4B). This effect was observed over a range of simulated cohesin off rates and topoisomerase I relaxation rates (fig. S5B).

To simulate the behavior of cohesin^EQ/EQ^ we modelled it as wild type cohesin, except that cohesin^EQ/EQ^ was only allowed to take a single loop-extrusion step, as experimentally observed (Janissen et al, 2024), whereupon cohesin^EQ/EQ^ stalled until it dissociated from DNA. Under these conditions, only negative supercoiling was predicted by the simulations at any cohesin-to-plasmid ratio, cohesin off rate, and topoisomerase I rate tested (Fig. 4C and fig. S5C), similar to our experimental observations (Fig. 1D and fig. S2G). This simulation result can be explained by the assumption made in our simulations that topoisomerase I can only alter Lk by multiples of 1 (ref. (*40*)). For this reason, topoisomerase I cannot resolve the -0.6 twist that is contained in the short loop that cohesin^EQ/EQ^ generates in a single extrusion step. In contrast, the unlooped region can accumulate twist or writhe from several cohesin^EQ/EQ^ complexes that results in accumulation of positive supercoils that can be resolved by topoisomerase I.

Since cohesin-SMC1^4E^ is predicted to bind DNA with reduced affinity, we reasoned that this mutant would be able to resist less torque in the clamped state when DNA becomes supercoiled. Indeed, when the stall torque, τ_stall_, of 12 k_B_T assumed for wild type cohesin (see Materials and Methods) was reduced to 8 k_B_T the cohesin-to-plasmid regime in which we observed negatively supercoiled plasmids vanished and instead positive supercoiling was predicted (Fig. 4D), in agreement with our experimental observations (Fig. 1F, 3B). With a limited number of simple and plausible assumptions, the simulations thus recapitulate the experimental results.

This indicates that the observed switch in supercoiling handedness is an indirect effect of the plasmid supercoiling assay, as opposed to cohesin itself changing the handedness of its supercoiling activity at high cohesin-to-plasmid ratios and when the DNA clamping sites are mutated. To understand this effect, we computed the fraction of plasmids containing negative or positive supercoiling at every time point and averaged the ratio of the positive to negative fraction. This revealed that plasmids accumulate on average more positively than negatively supercoiled regions at low wild type cohesin-to-plasmid ratios, but more negatively supercoiled regions at high ratios (Fig. 4F). In contrast, negatively supercoiled regions predominate at any ratio in plasmids exposed to cohesin-SMC1^4E^, and positively supercoiled regions in plasmids treated with cohesin^EQ/EQ^. If topoisomerase I can bind and relax all regions of the plasmid similarly well, as we assumed, this would explain why different concentrations and forms of cohesin would result in plasmids with different supercoiling handedness.

### Wild type cohesin and cohesin^EQ/EQ^ ‘protect’ supercoils, but cohesin-SMC1^4E^ does not

To explain why type IB topoisomerases such as human topoisomerase I do not resolve all supercoils in plasmids it was proposed that condensin protects the supercoils it has generated in the extruded DNA loop, so that topoisomerases can only resolve compensatory supercoils (*32, 33*). In contrast, our simulations predict the persistence of supercoils without assuming that cohesin protects supercoils, indicating that such a protection mechanism is not required to explain the supercoiling observed in assays containing cohesin or condensin.

However, our simulations do not exclude that cohesin protects supercoils. We therefore analyzed cohesin’s ability to induce supercoiling in the presence of TopA, which exclusively relaxes negative supercoils (ref. (*41*); fig. S6). If cohesin did *not* protect supercoils, TopA would resolve the negative supercoils induced by cohesin and should change the Lk of the plasmids. However, if cohesin did protect the negative supercoils it had induced, TopA should not be able to change Lk since the compensatory positive supercoils cannot be resolved by TopA. No ATP dependent supercoiling could be detected in the presence of low ratios of wild type cohesin and cohesin^EQ/EQ^ to plasmids, consistent with the possibility that these complexes *can* protect the negative supercoils they induced from TopA. Unexpectedly, however, positively supercoiled plasmids were detected after incubation with cohesin-SMC1^4E^ (fig. S6B), indicating that this mutant cohesin cannot protect the positive supercoils it has induced.

This finding could be explained if cohesin-SMC1^4E^ generated positively supercoiled DNA loops, an interpretation that would be consistent with all topoisomerase I and TopA plasmid supercoiling data (Fig. 1F, 3B, fig. S4, S5A and S6B) but not with our finding that cohesin-SMC1^4E^ generates negative twist in the magnetic tweezers assay (Fig. 3C–F). Another possibility is that the DNA within extruded loops might not be sufficiently negatively supercoiled to expose single-stranded DNA (ssDNA), a prerequisite for relaxation by TopA (ref. (*42*)). When we included this assumption in our simulations (see Methods) they indeed predicted no supercoiling by wild type cohesin or cohesin^EQ/EQ^ in the presence of TopA (fig. S6C and S6D), matching our experiments. However, the simulations also predicted no supercoiling in the presence of cohesin-SMC1^4E^ (fig. S6E), in contrast to our experimental results. These findings suggest that either cohesin-SMC1^4E^ exposes more ssDNA during supercoiling than wild type cohesin and cohesin^EQ/EQ^ or that cohesin-SMC1^4E^ differs in some yet unknown way from the other versions of cohesin.

### Testable predictions of the supercoiling-during-clamping hypothesis

Our results suggest that cohesin negatively supercoils DNA during clamping and that clamp mutations indirectly cause a change in supercoiling handedness in plasmid assays. This hypothesis makes several testable predictions. One of these is that weakening electrostatic interactions in the DNA clamp should also reduce supercoiling and induce a switch in supercoiling handedness. Remarkably, increased salt concentrations were indeed sufficient to generate positively supercoiled plasmids in the presence of wild type cohesin at low cohesin-to-plasmid ratios (Fig. 4G, compare lanes 2 and 14).

Cryo-EM and smFRET experiments indicate that NIPBL is required for clamping DNA (ref. (*12, 15-17*)) but have implied that the subunit STAG1 is not because clamping can be observed in the absence of this subunit (Scc3 in budding yeast; ref. (*15*)). The hypothesis that supercoiling occurs during DNA clamping therefore predicts that NIPBL is required for supercoiling, as we had observed (Fig. 1C), but that STAG1 might not be. Indeed, little difference in cohesin mediated supercoiling activity was observed in the absence and presence of STAG1 (Fig. 4H, lanes 6 and 8). A similar situation has been observed for cohesin’s ATPase activity, which also requires NIPBL-MAU2 but not STAG1 (*10, 43*). This similarity implies that cohesin’s DNA-dependent ATPase and supercoiling activities are linked, possibly because ATP-dependent DNA clamping is required for both.

Finally, if cohesin-SMC1^4E^ generates positively supercoiled plasmids by negatively supercoiling DNA, then limiting the activity of cohesin-SMC1^4E^ to one step should reveal its negative supercoiling activity. Indeed, introducing EQ/EQ mutations into cohesin-SMC1^4E^ reverted its positive supercoiling activity back to negative supercoiling (Fig. 4I). This epistatic effect was also predicted by our simulations (Fig. 4E). In contrast, introducing the SR/SR mutations into cohesin-SMC1^4E^ greatly reduced supercoiling, supporting the hypothesis that supercoiling occurs upon head engagement and DNA clamping (Fig. 4I).

### Cohesin-SMC1^4E^ forms shorter loops in cells

To test the effect of DNA clamping mutants on genome architecture in cells, we expressed either wild type SMC1 or SMC1^4E^ from a doxycycline inducible promoter in G1 synchronized HeLa cells, depleted endogenous SMC1 by auxin inducible degradation (*44*) and analyzed long-range chromosomal *cis*-interactions by Hi-C. In cells expressing cohesin-SMC1^4E^, long-range interactions were reduced genome wide (Fig. 5A, B) and long corner peaks (chromatin loops) were reduced, whereas short-range interactions and short corner peaks were increased (Fig. 5C), although SMC1^4E^ associated with chromatin similarly well as wild type SMC1 (fig. S7A). TAD numbers were not changed (fig. S7B), but interestingly the total number of corner peaks and TAD insulation were slightly increased in the presence of cohesin-SMC1^4E^ (Fig. 5D and fig.S7C).

**Fig. 5.**
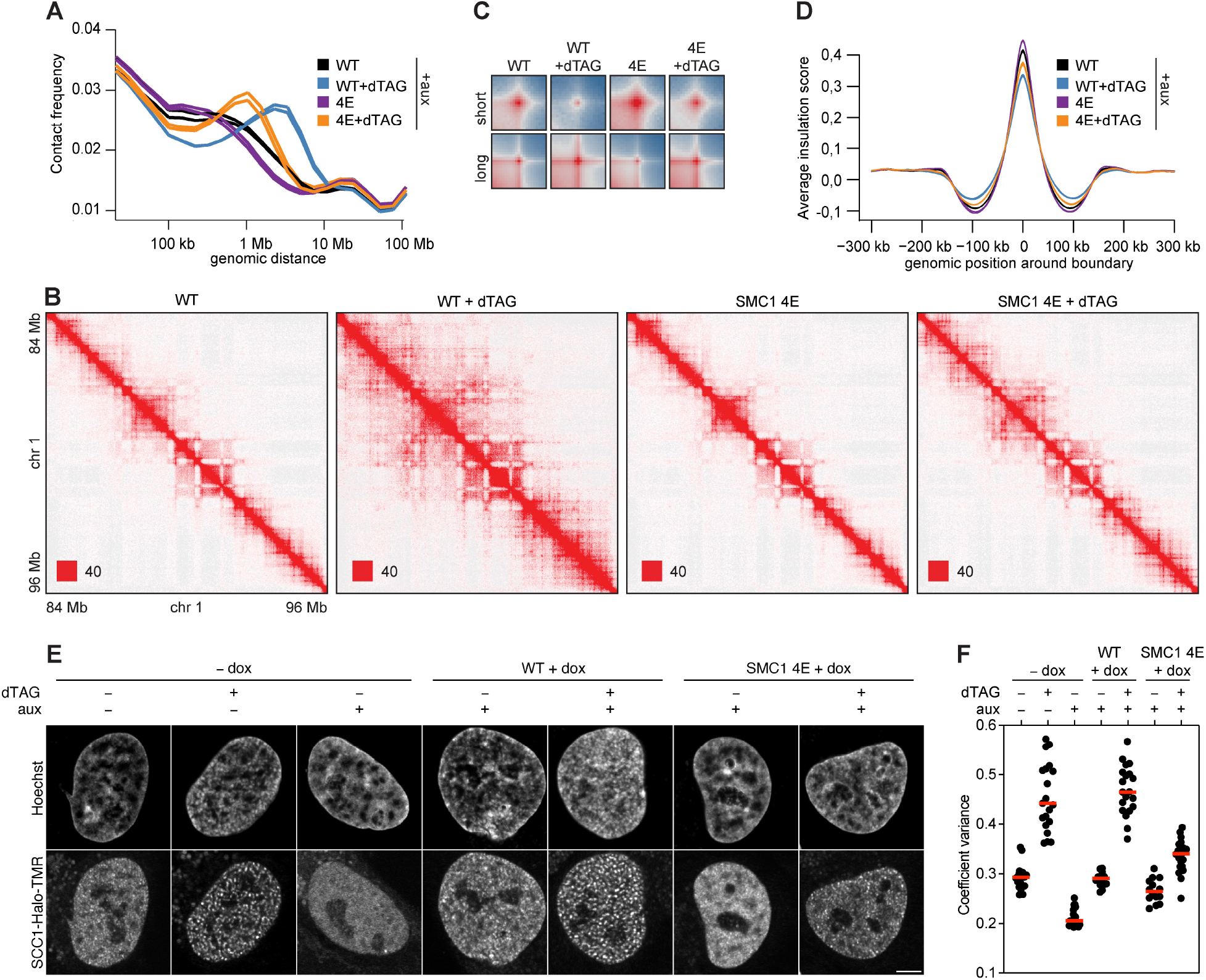
Cohesin-SMC1^4E^ forms shorter loops in cells. **(A)** Intra-chromosomal Hi-C contact frequency distribution plotted as a function of genomic distance in cells optionally treated with auxin, dTAG and doxycycline to degrade endogenous SMC1, degrade endogenous WAPL, and induce expression of wild type SMC1 or SMC1^4E^, respectively. **(B)** Balanced (Knight-Ruiz) normalized Hi-C contact matrices of chromosome 1 (84 - 96 Mb) from cells as described in (A). **(C)** Aggregated Hi-C peak analysis from cells as described in (A). Short: chromatin loops >100 kb at coordinates identified in SCC1-GFP HeLa cells. Long: chromatin loops > 500 kb at coordinates identified in auxin-treated WAPL-AID HeLa cells. **(D)** Average insulation score around TAD boundaries in cells as described in (A). **(E)** Representative live cell images of SCC1-HaloTMR in cells optionally treated with auxin, dTAG and doxycycline as described in (A). DNA was stained with Hoechst. Scale bars, 5 μm. **(F)** Quantification of vermicelli by measuring the coefficient variation of SCC1-HaloTMR fluorescence intensity (red bar denotes mean; data points are from individual cells obtained from three independent experiments).

To test whether these phenotypes are caused by increased sensitivity of cohesin-SMC1^4E^ to WAPL, we induced degradation of FKBP-tagged WAPL in these cells by dTAG. This increased SMC1^4E^ levels on chromatin (fig. S7A), long-range interactions (Fig. 5A), the length of corner peaks (Fig. 5C) and reduced TAD insulation (Fig. 5D), but not to the extent seen in the presence of wild type SMC1. These data indicate that cohesin-SMC1^4E^ is not hyper-sensitive to release by WAPL but instead causes the above Hi-C phenotypes by defects in DNA supercoiling and extrusion. Fluorescence microscopy of WAPL depleted cells expressing SMC1^4E^ revealed that much less cohesin accumulated in axial chromosomal domains called ‘vermicelli’ (ref. (*45*)) compared to cells expressing wild type SMC1 (Fig. 5E, F), further supporting the notion that cohesin-SMC1^4E^ is reduced in its ability to form long chromatin loops. These results suggest that cohesin’s negative supercoiling activity is required for DNA loop extrusion not only *in vitro* but also for chromatin looping in cells. These data also indicate that the strength of TAD boundaries depends on cohesin’s loop extruding activity, consistent with weakening of TAD boundaries in WAPL depleted cells (Fig. 5D; ref. (*4, 46*)).

## Discussion and conclusions

Our results show that cohesin supercoils DNA and indicate that this activity is an integral part of the loop-extrusion process. The accompanying manuscript by Janissen et al. demonstrates that negative supercoiling is a universal feature of all eukaryotic SMC complexes (Janissen et al., 2024), consistent with earlier observations that condensin supercoils DNA (*31-33*) and that oligonucleotides bound to the *E. coli* SMC complex MukBEF are oriented as if they represent parts of a negatively supercoiled loop (*47*). These findings provide fundamental insight into the mechanism of DNA loop extrusion, have important implications for the regulation of genome architecture and suggest that supercoiling is a common feature of SMC-mediated genome folding in all kingdoms of life.

Key steps in the loop extrusion cycle are the ATP-driven engagement of cohesin’s ATPase heads, resulting in ∼15 pN force generation (*18*), DNA translocation (∼100-200 bp/step (*19, 48*)) and subsequent DNA clamping. Our results indicate that cohesin supercoils DNA during these steps, implying that head engagement and force generation are not only used to translocate DNA but also to supercoil it. Our finding that DNA binding sites in the clamp and at the hinge are needed for supercoiling suggests that this process occurs when DNA is constrained between the hinge and the clamp, whereas a DNA binding site on STAG1 (*12, 49*) is dispensable for supercoiling and must therefore have another function during loop extrusion (*12*). Since cohesin’s coiled coils can twist around each other (*12*), it will be interesting to test whether supercoiling depends on or promotes these coiled coil movements.

Cohesin is sensitive to DNA tension, halting loop extrusion once a certain stall force has been reached (*19, 50*). Our results suggest that this sensitivity is not restricted to linear force, but that cohesin is also sensitive to torque in the DNA, and that cohesin’s stall torque depends on the DNA binding affinity of the clamp (fig. S7D). Our observation that cohesin mutated in one of the clamp’s DNA binding sites (*12*) and predicted to have a reduced stall torque (cohesin-SMC1^4E^) forms shorter genomic contacts, implies that cohesin’s ability to supercoil DNA is important for chromatin looping in cells. Cohesin-dependent genomic contacts detected by Hi-C might therefore represent both loops and supercoiled DNA, such as plectonemic structures.

The ability of the barrier protein CTCF to block loop extrusion increases with DNA tension (*19*). It is therefore conceivable that loop extrusion-induced torque at CTCF sites contributes to its barrier activity and may explain why Hi-C contacts are particularly enriched at these positions upon expression of SMC1^4E^ (fig. S7C).

Our findings also suggest that topoisomerases are required to prevent stalling of SMC complexes, which otherwise would stop extruding DNA once they had reached their stall torque, consistent with phenotypes obtained by topoisomerase inhibition (*51*). This hypothesis can explain why topoisomerase II accumulates at the base of cohesin and condensin loops (*20-23, 26-30*). Topoisomerases might therefore be particularly important for the formation of long loops, such as those that mediate protocadherin promoter choice (*5*) and immunoglobulin gene recombination (*6, 7*). Our results also provide evidence that cohesin’s *in vitro* loop extrusion activity has an important role in chromatin looping in cells, at variance with a recent proposal (*52*).

## Acknowledgments

We thank Daniela Goetz for cloning trimeric cohesin, the staff at IMBA/IMP/GMI BioOptics, Molecular Biology Service and Next-generation sequencing for technical support and all Peters and Dekker lab members for discussions. J-MP is also an adjunct professor at the Medical University of Vienna.

## Funding

European Research Council Advanced Grant 883684 (DNA looping) (CD)

NWO Grant OCENW.GROOT.2019.012 (CD)

BaSyC Program (CD) Boehringer Ingelheim (J-MP)

Austrian Life Sciences Programme 2023 (LS23 IF, project FO999902549) (J-MP)

European Research Council Horizon 2020 Research and Innovation Programme 101020558 (J-MP)

Human Frontier Science Program RGP0057/2018 (J-MP)

Vienna Science and Technology Fund LS19-029 (J-MP).

## Author contributions

IFD, SH, KN and J-MP designed experiments. IFD generated constructs for protein expression, purified proteins, performed plasmid supercoiling experiments and analyzed data. RB and RJ performed magnetic tweezers experiments. RJ analyzed magnetic tweezers data. RB performed simulations. SH performed high speed atomic force microscopy experiments and analyzed data. KN generated HeLa cell expression constructs and cell lines and performed and analyzed cellular experiments. GW purified human topoisomerase I and generated Hi-C libraries. RRS analyzed Hi-C data. BB generated cohesin ATPase head and hinge DNA binding mutant constructs for protein expression. IFD and J-MP wrote the manuscript with input from all authors. J-MP and CD supervised the study.

## Data and materials availability

All data are available in the main text or the supplementary materials. All plasmids used in this study are available from the lead contact with a completed materials transfer agreement.

## Supplementary Materials

Materials and Methods

Supplementary Text

Figs. S1 to S7

Tables S1 to S2

Movies S1 to S4

