## Supplementary Materials for "Cohesin supercoils DNA during loop extrusion"

**The PDF file includes:**

Materials and Methods
Supplementary Text
Figs. S1 to S7
Tables S1 to S2

**Other Supplementary Materials for this manuscript include the following:**

Movies S1 to S4

### Materials and Methods

#### Cloning and mutagenesis

A list of all constructs used in this manuscript is shown in table S2. Site-directed mutagenesis using Phusion Hot Start Flex DNA Polymerase (NEB) was used to introduce the individual '1E' mutations (K52E, R57E, K59E, K62E) into pLib SMC1, S1129R into pLib SMC1, S1116R into pLib SMC3-FLAG, and E1157Q and S1129R into pLib SMC1$^{4E}$. All other constructs produced in this study were generated using Gibson Assembly (*53*).

SMC1 and SMC3 expression cassettes from pLib constructs were combined into pBig1a polycistronic expression vectors using bigBAC (ref. (*54*)). SCC1 and STAG1 expression cassettes were combined into pBig1b vectors. pBig1a and pBig1b constructs were combined into pBig2ab vectors for wild type cohesin (table S2).

For the expression of wild type tetrameric cohesin (see below), Sf9 insect cells (Thermo Fisher Scientific; B82501) were either co-infected with baculoviruses generated from pBig1a SMC1_SMC3-FLAG and pBig1b SCC1$^{R172A/D279A/R450A}$-HALO_10xHIS-STAG1 vectors or infected with a baculovirus generated from a pBig2ab SMC1_SMC3-FLAG_ SCC1$^{R172A/D279A/R450A}$-HALO_10xHIS-STAG1 vector. All mutant forms of cohesin were generated by co-infection of two pBig1a and pBig1b baculoviruses. For trimeric cohesin, pBig1a SMC1, SMC3-FLAG and pBig1b HIS-SCC1(TEV)-HALO were combined into a pBig2ab vector.

#### Protein expression and purification

pLib, pBig1a, pBig1b and pBig2ab vectors were transformed into DH10EmBacY cells and bacmid DNA was isolated as described (*55*). Sf9 insect cells were transfected with bacmid DNA using Fugene 6 (Promega). The baculovirus-containing cell supernatant (V0) was collected 96 hours after transfection and used to infect 50 ml Sf9 cells at a density of 1 x$10^6$ cells/ml. Cells were grown in Grace medium at 100 rpm and 27 °C and centrifuged 72 hours after infection and the cell supernatant (V1) was then used to directly infect expression cultures at a density of 1.2 x$10^6$ cells/ml. Expression cultures were grown in Grace medium at 100 rpm and 27 °C. Cells were centrifuged around 54 hours after infection, washed in PBS, frozen in liquid nitrogen and stored at -80 °C.

<u>Cohesin and NIPBL-MAU2 purification</u>

Wild type and mutant forms of recombinant cohesin were purified from around 20 ml cell pellets as described (*10*). NIPBL-MAU2 was purified from around 20 ml cell pellets as described (*10*). STAG1 was purified from around 20 ml cell pellets as described (*12*).

<u>PDS5A-HALO-HIS purification</u>

All steps were performed at 4 °C unless indicated otherwise. Around 20 ml cell pellet was thawed and lysed by Dounce homogenization in PDS5 purification buffer (25 mM $Na_2HPO_4/NaH_2PO_4$ [pH 7.5], 500 mM NaCl, 5 % glycerol) supplemented with 10 mM imidazole [pH 7.5], 0.05 % Tween 20, 1 mM PMSF, 3 mM betamercaptoethanol, 10 µg/ml aprotinin, 2 mM benzamidine and cOmplete EDTA-free protease inhibitor cocktail (Merck; 11873580001). After centrifugation (47000 g, 45 min), the soluble fraction was combined with 1 ml of NiNTA agarose (Qiagen; 30230) and incubated for 90 min. Beads were washed with 3 x 20 bead volumes of PDS5 purification buffer supplemented with 30 mM imidazole [pH 7.5] and 3 mM betamercaptoethanol. Bound protein was eluted with 6 ml PDS5 elution buffer (25 mM $Na_2HPO_4/NaH_2PO_4$ [pH 7.5], 150 mM NaCl, 5 % glycerol, 300 mM imidazole [pH 7.5], 1 mM DTT). Eluted protein was concentrated to around 0.5 ml using Vivaspin 6 50 kDa MWCO ultrafiltration units (Sartorius; VS0631), filtered and applied to a Superdex 200 10/300 GL column at 0.4 ml/min equilibrated in size exclusion buffer (25 mM $Na_2HPO_4/NaH_2PO_4$ [pH 7.5], 150 mM NaCl, 5 % glycerol, 1 mM DTT). Fractions containing PDS5A-HALO-HIS were pooled, concentrated, frozen in liquid nitrogen and stored at -80°C.

Human Topoisomerase I purification

All steps were performed at 4 °C unless indicated otherwise. Around 20 ml cell pellet was thawed and lysed by Dounce homogenization in TopoI purification buffer 1 (25 mM $Na_2HPO_4/NaH_2PO_4$ [pH 7.5], 500 mM NaCl, 5 % glycerol) supplemented with 10 mM imidazole [pH 7.5], 0.05 % Tween 20, 1 mM PMSF, 3 mM betamercaptoethanol, 10 μg/ml aprotinin, 2 mM benzamidine and cOmplete EDTA-free protease inhibitor cocktail (Merck; 11873580001). After centrifugation (47000 g, 45 min), the soluble fraction was combined with 4 ml of Toyopearl AF-chelate-650M resin (Tosoh Bioscience) pre-charged with Ni2+ ions and incubated for 2 h. Beads were washed with 3 x 10 bead volumes of TopoI purification buffer 1 supplemented with 25 mM imidazole [pH 7.5]. Bound protein was eluted with 25 ml TopoI purification buffer 2 (25 mM $Na_2HPO_4/NaH_2PO_4$ [pH 7.5], 150 mM NaCl, 5 % glycerol, 300 mM imidazole [pH 7.5]). The eluate was then combined with 5 ml of FLAG-M2 agarose resin (Sigma; A2220) and incubated for three hours. Beads were washed with 3 x 10 bead volumes of TopoI purification buffer 3 (25 mM $Na_2HPO_4/NaH_2PO_4$ pH 7.5, 150 mM NaCl, 5 % glycerol, 50 mM imidazole pH 7.5). Bound protein was eluted with 15 ml TopoI purification buffer 3 supplemented with 0.5 mg/ml 3xFlag peptide. Eluates were concentrated to ~0.75 ml using Vivaspin 20 100 kDa MWCO ultrafiltration units (Sartorius; VS2042), frozen in liquid nitrogen and stored at -80°C.

**Human topoisomerase I DNA supercoiling assay**

Negatively supercoiled plasmid DNA (pH6HTN (Promega), 4 kb) was relaxed by incubating it together with human topoisomerase I for 45 min at 37 ºC. This mixture was then supplemented with recombinant human cohesin and NIPBL-MAU2 as indicated and incubated for a further 90 min (final reaction conditions: 20 mM $Na_2HPO_4/NaH_2PO_4$ [pH 7.5], 5 mM Tris [pH 7.5], 25 mM NaCl, 2.5 mM $MgCl_2$, 1 mM DTT, 0.1 mg/ml BSA (Thermo Fisher Scientific; AM2616), 2 mM ATP (Jena Biosciences; NU-1049), 0.4 nM pH6HTN, 3 nM human topoisomerase I, 0 - 32 nM cohesin (corresponding to a cohesin:plasmid molar ratio of up to 80:1) and 0 – 64 nM NIPBL-MAU2 (i.e. a two-fold molar excess relative to cohesin) in a 100 μl reaction). Reactions were stopped by addition of sodium dodecyl sulphate (SDS; 0.33 % final), EDTA (6.7 mM final) and Proteinase K (0.1 mg/ml final; Sigma-Aldrich; P6556) and incubated for 40 min at 50 °C. DNA was extracted using phenol:chloroform:isoamyl alcohol (Sigma-Aldrich; P2069) and then chloroform (Sigma-Aldrich; P32211) in 5PRIME Phase Lock Gel Light tubes (Quantabio; 2302820). Purified DNA was supplemented with glycerol to 10 % final concentration and trace amounts of Orange G (Sigma-Aldrich; P2069). Half of the reaction volume were electrophoresed in a 0.7 % agarose gel in 1x TBE buffer supplemented with 0.4 μg/ml chloroquine (Sigma-Aldrich; C6628) for 17 hours at 32 V. Gels were stained with Sybr Gold (Thermo Fisher Scientific; S11494) and visualized using a ChemiDoc MP imager (Bio-Rad).

Human topoisomerase I DNA supercoiling assay using pre-relaxed plasmid DNA as input

For the experiment described in fig. S2A and fig. S2B in which plasmid DNA was relaxed and purified prior to incubation with cohesin, negatively supercoiled plasmid DNA (pH6HTN) was relaxed by incubating it together with human topoisomerase I for 45 min at 37 °C (final reaction conditions: 20 mM $Na_2HPO_4/NaH_2PO_4$ [pH 7.5], 5 mM Tris [pH 7.5], 25 mM NaCl, 2.5 mM $MgCl_2$, 1 mM DTT, 0.1 mg/ml BSA (Thermo Fisher Scientific; AM2616), 0.4 nM pH6HTN, 3 nM human topoisomerase I in a 600 µl reaction). Reactions were stopped and purified as above and further purified using Ampure XP beads according to the manufacturer's instructions (Beckmann). This relaxed DNA was then incubated in the presence or absence of human topoisomerase I, cohesin and NIPBL-MAU2 as described above.

Determination of cohesin-mediated supercoiling handedness using Topo IV, Gyrase and TopA

For the experiments described in fig. S4A–D, negatively supercoiled plasmid DNA (pH6HTN) was relaxed by incubating it together with human topoisomerase I for 45 min at 37°C. This mixture was then supplemented with cohesin$^{EQ/EQ}$ or cohesin SMC1$^{4E}$ and NIPBL-MAU2 as indicated and incubated for a further 90 min (final reaction conditions: 20 mM $Na_2HPO_4/NaH_2PO_4$ [pH 7.5], 5 mM Tris [pH 7.5], 25 mM NaCl, 2.5 mM $MgCl_2$, 1 mM DTT, 0.1 mg/ml BSA (Thermo Fisher Scientific; AM2616), 2 mM ATP (Jena Biosciences; NU-1049), 0.4 nM pH6HTN, 3 nM human topoisomerase I, 6 nM cohesin (corresponding to a cohesin:plasmid molar ratio of 15:1) and 12 nM NIPBL-MAU2 (i.e. a two-fold molar excess relative to cohesin) in a 600 µl reaction). Reactions were stopped and purified as above and further purified using Ampure XP beads according to the manufacturer's instructions (Beckmann). 50 ng of this 'EQ/EQ DNA' or 'SMC1$^{4E}$ DNA' was then incubated with DNA gyrase (Inspiralis; 0 – 2 units), *E. coli* Topo IV (Inspiralis; 0 – 0.05 units) or *E. coli* TopA (New England Biolabs; 0 – 5 units) for 40 min at 37 °C according to manufacturer's instructions. Reactions were stopped, purified and electrophoresed as above. See also Supplementary Text.

For the experiment described in Figure S4E, 50 ng of negatively supercoiled plasmid DNA (pH6HTN (Promega), 4 kb) was incubated with *E. coli* TopA (New England Biolabs; 0 – 5 units) for 40 min at 37 °C according to manufacturer's instructions. Reactions were stopped, purified and electrophoresed as above. See also Supplementary Text.

Comparison between cohesin-mediated supercoiling in the presence of human topoisomerase I and E. coli TopA

For the experiment described in fig. S6A – B in which supercoiling by cohesin in the presence of human topoisomerase I and *E. coli* TopA was compared, negatively supercoiled plasmid DNA (pH6HTN) was relaxed by incubating it together with human topoisomerase I or *E. coli* TopA for 45 min or 20 min, respectively, at 37°C. These mixtures was then supplemented with recombinant human cohesin and NIPBL-MAU2 as indicated and incubated for a further 90 min (final reaction conditions: 20 mM $Na_2HPO_4/NaH_2PO_4$ [pH 7.5], 5 mM Tris [pH 7.5], 25 mM NaCl, 2.5 mM $MgCl_2$, 1 mM DTT, 0.1 mg/ml BSA (Thermo Fisher Scientific; AM2616), 2 mM ATP (Jena Biosciences; NU-1049), 0.4 nM pH6HTN, 6 nM cohesin (corresponding to a cohesin:plasmid molar ratio of 15:1), 12 nM NIPBL-MAU2 (i.e. a two-fold molar excess relative to cohesin) and either 3 nM human topoisomerase I or 2.5 units *E. coli* TopA in a 100 µl reaction). Reactions were stopped, purified and electrophoresed as described in the human topoisomerase I DNA supercoiling assay section.

**Quantification of topoisomer distributions in chloroquine gels**

To determine the linking number change between conditions, gels were quantified according to (*33, 56*). Briefly, the relative intensity of individual topoisomers per condition was determined using ImageJ (NIH). The mean of each distribution (Lk) was determined by Gaussian fitting in GraphPad Prism. ΔLkATP for the indicated form of cohesin was calculated by subtracting the Lk values obtained in the presence and absence of ATP. To estimate the ATP-dependent change in linking number per cohesin, ΔLkATP was divided by the cohesin:plasmid ratio.

**High-speed Atomic Force Microscopy**

Negatively supercoiled plasmid DNA (pH6HTN) was relaxed by incubating it together with human topoisomerase I for 45 min at 37°C. This mixture was then supplemented with recombinant human cohesin and NIPBL-MAU2 and incubated for a further 90 min (final reaction conditions: 20 mM $Na_2HPO_4$/$NaH_2PO_4$ [pH 7.5], 5 mM Tris [pH 7.5], 25 mM NaCl, 2.5 mM $MgCl_2$, 1 mM DTT, 0.1 mg/ml BSA (Thermo Fisher Scientific; AM2616), 2 mM ATP (Jena Biosciences; NU-1049), 1.2 nM pH6HTN, 9 nM human topoisomerase I, 18 nM cohesin (corresponding to a cohesin:plasmid molar ratio of 15:1) and 32 nM NIPBL-MAU2 in a 100 µl reaction). Reactions were stopped and purified as described in the human topoisomerase I DNA supercoiling assay section.

HS-AFM measurements were performed as previously described for dynamic molecules (*57*). In brief, a mica sheet was glued to a glass rod (1.5 mm in diameter, 2 mm height, Hilgenberg GmbH) using two-compound epoxy glue (UHU, Bolton Adhesives). The opposite end of the glass rod was attached to the Z scanner using wax.

Purified DNA samples from were deposited onto a freshly cleaved mica sheet and incubated for 5 min at room temperature. To remove unbound DNA plasmids, the surface was rinsed 5 times with 2 µL imaging buffer (10 mM Tris [pH 8.0], 1 mM EDTA, 12.5 mM $MgCl_2$). DNA plasmids were then visualized in imaging buffer with a scanning HS-AFM (RIBM) operated in tapping mode. The free amplitude was set to 1 nm and the amplitude setpoint to 90 % of the free oscillation amplitude. Ultrashort cantilevers (USC-F1.2-k0.15, NanoWorld, Switzerland) with a spring constant of 0.15 N/m, a resonance frequency of around 0.6 MHz and a quality factor of around 2 in liquid were used. Areas of 640 nm x 640 nm were imaged at 500 ms/frame, which typically allowed visualization of one or two DNA plasmids. Each plasmid was imaged for 60 frames and the number of DNA crossings was counted. All HS-AFM measurements were performed at room temperature.

HS-AFM data was collected using Igor Pro software (Wave Metrics Inc., Lake Oswego, OR, USA). Images were processed by applying mean-flatten and resonance noise correction filters using Kodec software version 4.3.6.16 (ref. (*58*)).

**Measurement of DNA loop extrusion-induced DNA twist using magnetic tweezers**

Magnetic tweezers experiments were performed using a 3.6 kbp torsionally constrained DNA as described (Janissen et al 2024). The cohesin flow-in mixture contained: 50 mM Tris [pH 7.5], 40 mM NaCl, 2.5 mM $MgCl_2$, 0.25 mg/ml BSA, 1 mM DTT, 0.05 % Tween20, 1 mM ATP, 20 pM cohesin $SMC1^{4E}$, 50 pM NIPBL-MAU2, 1 mM ATP.

**1D cohesin supercoiling simulations**

We use a fixed-time-step Monte Carlo algorithm for 1D simulations. Each lattice sites corresponds to 1 bp and the plasmid was defined with a length of L=4000 bp, with periodic boundary conditions (lattice site 4000 equals lattice site 0). Simulations were carried out at 297 K ($k_BT$=4.1 pN nm). A time step equals one loop extrusion step, which roughly translates to 0.5 s, based on the ATPase activity of cohesin of roughly 2/protein/s (ref. (*10*)). The simulation was run for 1800 time steps (~ 15 min).

For every time step, the following modules are executed:

Cohesin complexes spawn
Cohesin lifetime and direction lifetime assignment
Cohesin stepping
Linking number fixation by topoisomerase
Cohesin dissociation

Cohesin complexes spawn

Cohesin complexes have a uniform binding probability in the absence of any accumulated twist (i.e. for the first cohesin complex binding to the plasmid). If twist has been accumulated in the plasmid, the cohesin complex binds to a positively supercoiled segment with a 79 % probability (*31*). The binding rate of cohesin to the plasmid, $k_{on}$, was varied over several orders of magnitude between 1 x$10^{-5}$ and 1 x$10^{2}$ and the number of cohesin complexes on the plasmid was counted during the simulation, which was used to evaluate the cohesin:plasmid ratio in simulations and to compare to experiments (e.g. fig. S5B).

Cohesin lifetime and direction lifetime assignment

Upon binding, the lifetime of the cohesin complex is drawn from an exponential distribution with mean $1/k_{off}$. Furthermore, cohesin complexes move directionally and can switch directions with rate constant $k_{switch}$=0.025 $s^{-1}$ (roughly once or twice per minute) and direction lifetimes are drawn from an exponential distribution with mean $1/k_{switch}$.

Cohesin stepping

All plasmid-bound cohesin complexes attempt to make a step in random order during one time step of the simulation. Each cohesin takes a step of $\Delta l$= 150 bp (ref. (*50*)), enlarging its loop, while a twist of $\Delta Tw$= -0.6 is added to the loop. If the segment on which the step is being made has a non-zero twist itself, a part of this twist is transferred into the loop. The relative twist per bp in a segment s is $rTw_s=Tw_s/L_s$, where $Tw_s$ is the accumulated twist and $L_s$ is the length of segment s. The twist transferred into the loop while the loop s is being enlarged on the expense of segment s is $\Delta Tw=-0.6+\Delta L\cdot rTw_s$. Concomitantly, a twist of $\Delta Tw=+0.6-\Delta L\cdot rTw_s$ is added to segment s (illustrated in Fig. 4A). Note that region s may also be a previously extruded loop such that the loading of the next SMC produces nested loops. Cohesin complexes are allowed to traverse one another (*39*) and generate Z-loops while doing so. If Z-loops are generated, twist is added to the resulting Z loop delimited by the two loop anchors defining the Z-loop. Similarly, Z loops can be resolved by nesting loops. The loop anchors are modelled as twist diffusion barriers.

After every potential move, the resulting torque τ in the loop as well as in segment s is computed via (*59*) $\tau=2\pi k_B T C \cdot Tw/L_c$, where C=100 nm is the twist persistence length of dsDNA (ref. (*60*)) and $L_c=L\cdot$ 0.342 nm/bp denotes the contour length of the plasmid. If the torque in all concerning segments remains below the stall torque, $\tau_{stall}$, the move is accepted and rejected otherwise. Even though not measured, cohesin complexes must have a stall torque which is lower than the energy of ATP binding to the complex. Here we set $\tau_{stall}$= 12 $k_BT$, slightly above the energetic cost to extrude DNA (*19, 61*). Lower values were used to simulate cohesin $SMC1^{4E}$ (Fig. 4D and 4E). $Cohesin^{EQ/EQ}$ or $cohesin^{EQ/EQ}$-$SMC1^{4E}$ was simulated by allowing only a single step before stalling (Fig. 4C and 4E).

Linking number fixation by topoisomerases

Topoisomerases bind to the plasmid at a rate $k_{topo}$. The binding position is drawn from a uniform distribution across the plasmid. Human topoisomerase I is able to relax both positive as well as negative supercoils up to a maximum of $\Delta L_k$=27 (ref. (*62*)), or all linking numbers if the segment topoisomerase bound to contains fewer linking numbers.
*E.coli* topoisomerase I (TopA) selectively relaxes only negative supercoils by recognizing ssDNA generated by underwinding of DNA (*42, 63*) and changes the linking number by exactly +1 (because it is a type IA topoisomerase that uses a strand passage mechanism; ref. (*64*)). In the absence of tension on the DNA, this occurs at a supercoiling density $\sigma=(L_k-L_{k0})/(L_{k0})$ ~-0.065 (ref. (*65*)), within the range of supercoiling density values found *in vivo* (*66, 67*). Simulations of experiments in the presence of TopA were thus carried out by allowing the topoisomerase to only act on segments in which $\sigma \leq -0.065$ (see main text). The results were consistent within the tested range $-0.075 \leq \sigma \leq -0.05$.

Cohesin dissociation

When a cohesin complex lifetime reaches zero, it dissociates from the plasmid. If a Z-loop is associated with the segment, this Z-loop is first resolved upon which the loop dissolves. The loop length and its twist merges with its surround segment whose relative twist and torque is updated.

Sensitivity to parameter choices

We varied the dissociation and the topoisomerase action rate, $k_{off}$ and $k_{topo}$, respectively. The results are qualitatively similar as long as the topoisomerase activity is not too low (e.g. $k_{topo} = 10^{-3}$/step causes only 1-2 topoisomerase actions per simulation run yielding only low amounts of fixed linking numbers (fig. S5B and fig. S5C).

**HeLa cell experiments**

HeLa cell line generation

Cells were cultured as described (*44*). SCC1-Halo-P2A-Tir1, SMC1-AID-mKate2, and Blasticidin-P2A-FKBPF36V-WAPL-expressing cells (*44*) were infected with lentiviruses to generate wildtype SMC1 or $SMC1^{4E}$-expressing cells as described (*44*).

Cell synchronization, chromatin fractionation and Hi-C library preparation

Cells were synchronized in G1 cell cycle stage using two consecutive rounds of treatment with 2 mM thymidine (Sigma-Aldrich) in the presence of 1 μg/ml doxycycline. Two hours prior to release from the second thymidine arrest, cells were treated with 200 μM auxin (Andole-3-

Acetic Acid, Gold Biotechnology) and 1 μM dTAG7 (TOCRIS Bioscience; 6912) . After washing with pre-warmed medium, cells were incubated with medium containing 200 μM auxin and 1 μM dTAG7 for 6 hours and harvested for chromatin fractionation and Hi-C. Chromatin fractionation and Hi-C library preparation were performed as described (*44*). Western blotting was performed using antibodies against SMC1 (in-house; A1027), SMC3 (Thermo Fisher Scientific; A300-060A; RRID: AB_67579), WAPL (in-house; A1017), Tubulin (Sigma-Aldrich; T5168; RRID: AB_477579) and Histone H3 (Cell Signalling Technology; 9715L; RRID: AB_331563).

<u>Hi-C data processing</u>

Hi-C data processing was performed as described (*44*).

<u>Live cell imaging and vermicelli quantification</u>

Cells were seeded on chambered coverglass (Nunc 155409) for 2 days in DMEM supplemented with 10% FCS, 0.2 mM L-glutamine, Penicillin-Streptomycin (Sigma-Aldrich; P0781) and 1 μg/ml doxycycline to induce expression of wild type SMC1 or $SMC1^{4E}$. Media was then supplemented with dTAG7 (1 μM) and cells were incubated for a further 5 hours. Before imaging SCC1-Halo, cells were incubated with 250 μM HaloTag TMR ligand for 20 min. After washing with pre-warmed cell culture medium three times, cells were incubated for 30 min and then exchanged for pre-warmed phenol red free medium supplemented with 10 % FCS, 0.2 mM L-glutamine and Penicillin-Streptomycin for imaging. For DNA labeling, cells were incubated with 0.1 μg/ml Hoechst 33342 for 20 min before imaging. Live cell imaging was performed using an LSM880 confocal microscope (Carl Zeiss), equipped with a 60x /1.4 numerical aperture (N/A) oil DIC Plan-Apochromat objective at 37 ºC and 5 % $CO_2$. Vermicelli were quantified as described (*44*).

**Supplementary Text**

<u>Determination of cohesin-mediated supercoiling handedness using Topo IV, Gyrase and TopA</u>

In chloroquine gel electrophoresis, negatively supercoiled plasmids can migrate faster than relaxed plasmids if the number of negative supercoils greatly exceeds the number of positive supercoils induced by chloroquine. This can be seen when negatively supercoiled plasmids isolated from *E. coli* were treated with TopA, which can only remove negative supercoils (*41*) (fig. S4E). Without TopA treatment, the electrophoretic mobility of these plasmids was high, indicating that they contained more negative supercoils than chloroquine-induced positive supercoils (fig. S4E, lane 2). TopA treatment reduced the mobility of these plasmids until the number of negative supercoils equaled the number of chloroquine-induced positive supercoils (fig. S4E, lane 4). Beyond this point, chloroquine-induced positive supercoils prevailed and increased the mobility of the plasmids again (fig. S4E, lane 5). These results illustrate that the supercoiling handedness of plasmids that migrate faster than relaxed plasmids in chloroquine gels cannot be unequivocally determined from the electrophoretic mobility of these DNA molecules alone.

We therefore tested whether the plasmids generated in the presence of cohesin-$SMC1^{4E}$ were indeed positively supercoiled, as we assumed, as opposed to representing highly negatively supercoiled plasmids. To do this, we performed human topoisomerase I DNA supercoiling assays in the presence of $cohesin^{EQ/EQ}$ or cohesin-$SMC1^{4E}$, purified this DNA and then incubated it with *E. coli* topoisomerase IV, gyrase or TopA, enzymes that differentially affect negatively and positively supercoiled plasmids (fig. S4A).

Topo IV relaxed supercoils generated by cohesin-SMC1$^{4E}$ at much lower concentrations than supercoils generated by cohesin$^{EQ/EQ}$ (fig. S4B). Because Topo IV preferentially (though not exclusively) relaxes positively supercoiled DNA (ref. (*68-70*)), these results suggest that plasmids indeed become negatively and positively supercoiled in the presence of cohesin$^{EQ/EQ}$ and cohesin-SMC1$^{4E}$, respectively.

This conclusion is supported by the effects we observed following treatment with DNA gyrase, which does not resolve but instead introduces negative supercoils (*71*). When incubated with plasmids supercoiled by cohesin$^{EQ/EQ}$, high concentrations of gyrase converted all slowly migrating DNA bands into one high-mobility species (fig. S4C, lane 6). This effect suggests that plasmids had become negatively supercoiled in the presence of cohesin$^{EQ/EQ}$ and were more negatively supercoiled by gyrase, further increasing their mobility. However, plasmids supercoiled by cohesin-SMC1$^{4E}$ showed a different response to increasing gyrase concentrations (fig. S4C, lanes 7 – 11). In this case, the fast-migrating plasmids generated in the presence of cohesin-SMC1$^{4E}$ were converted into slower migrating bands at the second highest gyrase concentration but re-converted into a distinct high-mobility species at the highest gyrase dose. This behavior suggests that plasmids had indeed become positively supercoiled in the presence of cohesin-SMC1$^{4E}$ and were gradually converted into negatively supercoiled forms at increasing gyrase concentrations.

The conclusion that plasmids become differently supercoiled in our assays is also supported by the effects that we observed with TopA. This enzyme could only relax plasmids previously incubated with cohesin$^{EQ/EQ}$ (fig. S4D) or negatively supercoiled plasmids isolated from *E. coli* (fig. S4E) or but not plasmids previously incubated with cohesin-SMC1$^{4E}$ (fig. S4D). Because TopA can only relax negatively supercoiled DNA (ref. (*41*)) these results indicate that plasmids had become negatively supercoiled in the presence of cohesin$^{EQ/EQ}$, but positively supercoiled in the presence of cohesin-SMC1$^{4E}$.

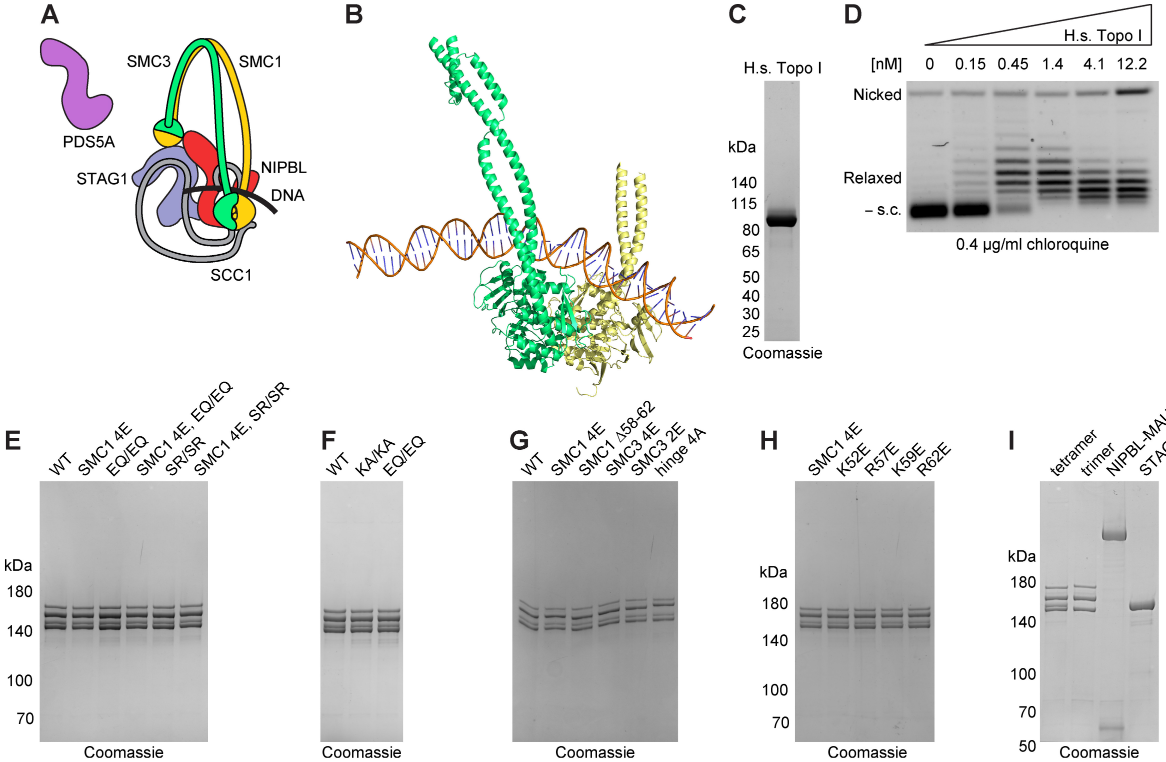


**Fig. S1. Characterization of recombinant human topoisomerase I and cohesin. (A)** Illustration of cohesin, NIPBL and PDS5A. Cohesin and NIPBL are represented in the DNA clamp state. **(B)** Cartoon representation of the ATPase head domains of SMC1 and SMC3 and DNA from the cryo-EM structure of human cohesin in the DNA clamp state (PDB: 6WG3). SMC1, SMC3 and DNA are colored yellow, green and orange, respectively. Cartoon representations of NIPBL, SCC1, STAG1 and the SMC1/SMC3 hinge domain are not displayed. **(C)** Coomassie staining of recombinant human topoisomerase I after SDS-PAGE. **(D)** Relaxation of negatively supercoiled plasmid DNA in the presence of increasing concentrations of human topoisomerase I. Representative image from 2 independent experiments. **(E – I)** Coomassie staining of the indicated forms of recombinant human cohesin, NIPBL-MAU2 and STAG1.

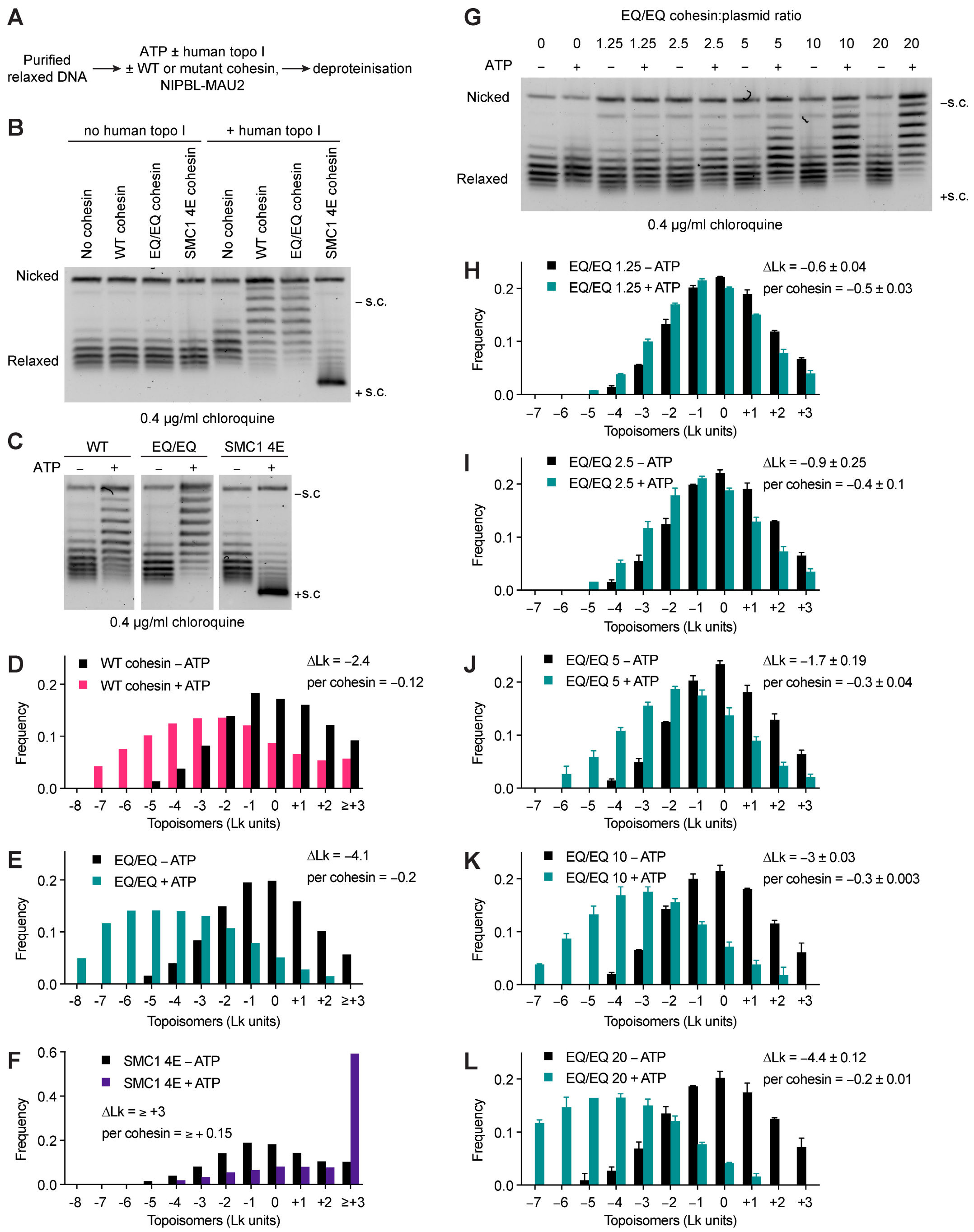


**Fig. S2. Characterization of cohesin-mediated supercoiling. (A)** Outline for experiment shown in panel (B). **(B)** Human topoisomerase I DNA supercoiling assay in the presence or absence of the indicated proteins using purified relaxed DNA as input. Cohesin was added at a 15:1 molar ratio relative to plasmid DNA. NIPBL-MAU2 was added to all reactions at a 30:1 molar ratio relative to plasmid DNA. Representative image from 2 independent experiments. **(C)** Human topoisomerase I DNA supercoiling assay in the presence or absence of the indicated proteins and ATP. Cohesin was added at a 20:1 molar ratio relative to plasmid DNA. NIPBL-MAU2 was added to all reactions at a 40:1 molar ratio relative to plasmid DNA. Representative image from 2 independent experiments. **(D – F)** To estimate ΔLk per cohesin molecule, we quantified the relative distribution of plasmid topoisomers in the presence and absence of ATP in the agarose gels shown in (C), and divided the difference between the two means (ΔLkATP) by the cohesin to plasmid ratio. **(G)** As (C) except cohesin$^{EQ/EQ}$ was added at the indicated cohesin:plasmid ratio along with NIPBL-MAU2 at a 40:1 NIPBL:plasmid ratio. Representative image from 2 independent experiments. **(H – L)** Relative intensity of topoisomer distributions quantified from the agarose gel shown in (G) and a replicate. ΔLk denotes the difference in Lk units between the midpoints of each data set. Error bars denote standard deviation.

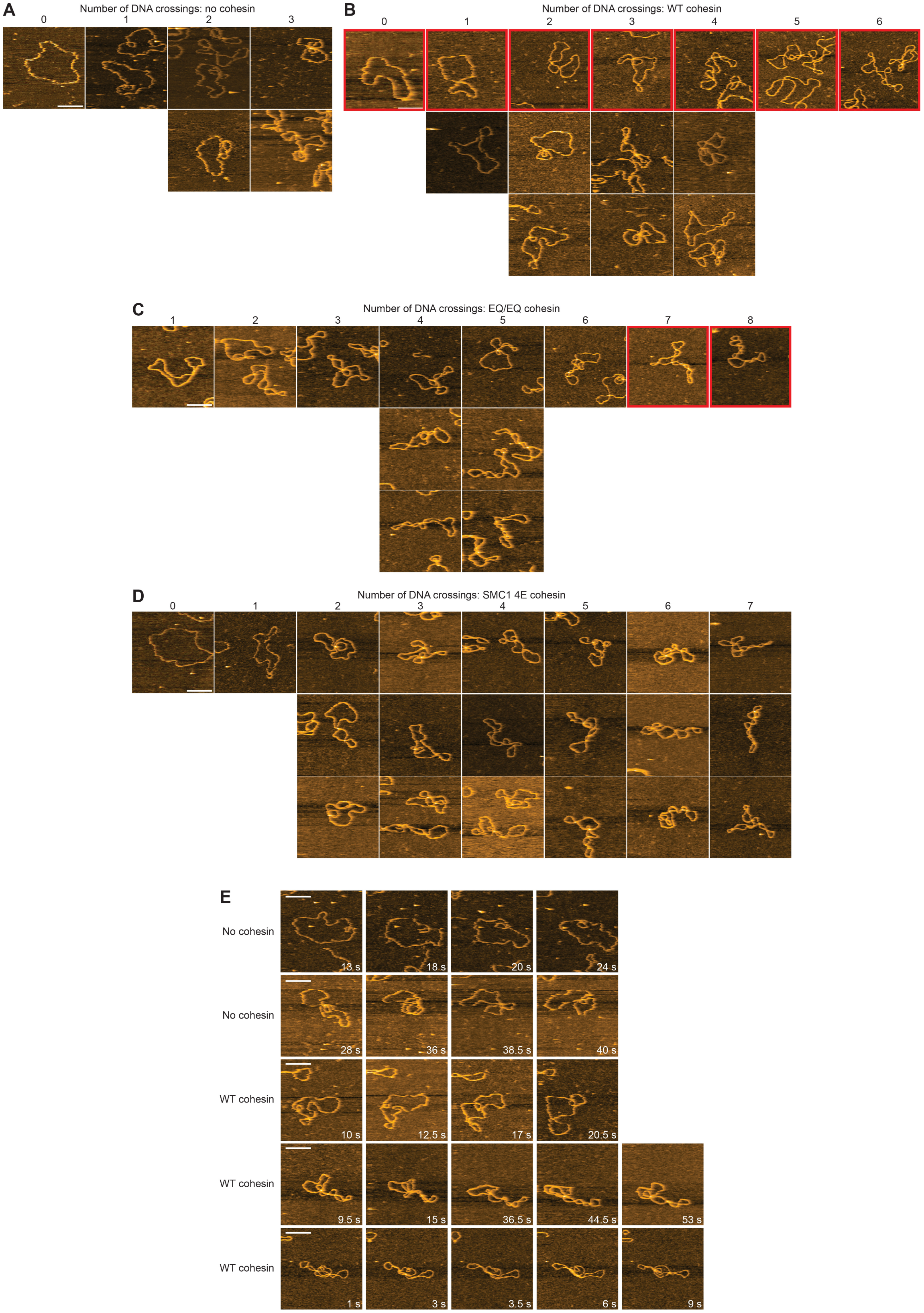


**Fig. S3. Additional representative images from HS-AFM recordings of the indicated forms of plasmid DNA. (A – D)** The red outlined images denote those used in Figure 2. **(E)** Timelapse images showing translocation of DNA crossings within plasmids previously incubated with human topoisomerase I and ATP in the presence or absence of wild type cohesin (15:1 cohesin:plasmid ratio) and NIPBL-MAU2 (30:1).

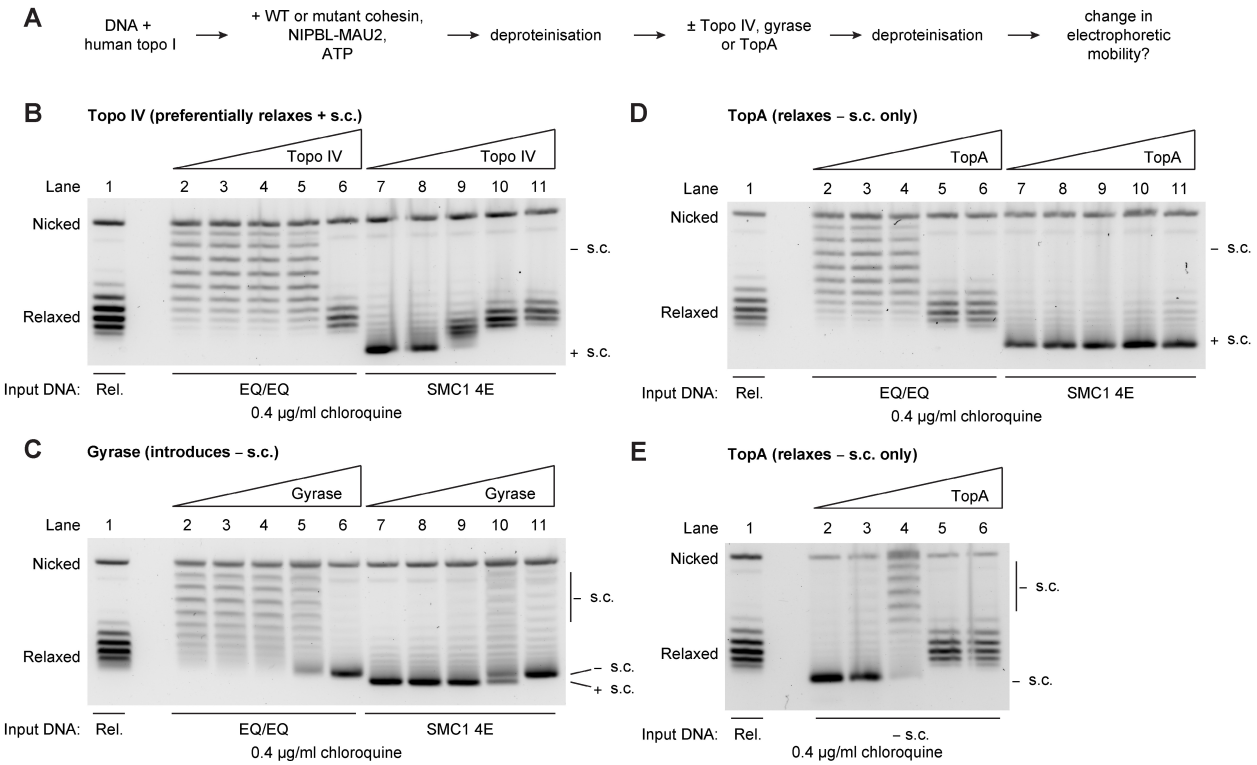


**Fig. S4. Cohesin$^{EQ/EQ}$ and cohesin-SMC1$^{4E}$ induce net negative and positive DNA supercoiling in the plasmid assay, respectively. (A)** Experiment outline for panels B – E. **(B)** Purified DNA previously incubated with human topoisomerase I and cohesin$^{EQ/EQ}$ or cohesin-SMC1$^{4E}$ was further incubated with increasing concentrations of *E. coli* Topo IV (0, 0.002, 0.006, 0.02, 0.05 units), which preferentially relaxes positively supercoiled DNA. Purified DNA previously incubated with human topoisomerase I was used as a marker for relaxed DNA. Higher concentrations of Topo IV were required to relax EQ/EQ DNA than SMC1 4E DNA, consistent with the hypothesis that cohesin$^{EQ/EQ}$ and cohesin-SMC1$^{4E}$ generated negatively and positively supercoiled plasmids in the presence of topoisomerase I, respectively. Representative image from 2 independent experiments. **(C)** As (B), except DNA was incubated with increasing concentrations of *E. coli* gyrase (0, 0.07, 0.22, 0.67, 2 units), which introduces negative DNA supercoils. At high concentrations, gyrase introduced sufficient negative supercoiling that EQ/EQ and SMC1$^{4E}$ DNA migrated as single bands (lanes 6 and 11). These highly negatively supercoiled plasmids migrated slower than the positively supercoiled plasmids observed following incubation with cohesin-SMC1$^{4E}$ in the absence of gyrase (lane 7) due to the differential effect of chloroquine-induced positive supercoiling on negatively or positively supercoiled DNA, respectively. Representative image from 2 independent experiments. **(D)** As (B) except DNA was incubated with increasing concentrations of *E. coli* TopA (0, 0.28, 0.83, 2.5, 5 units), which relaxes negative DNA supercoils only. TopA relaxed EQ/EQ DNA but did not change the mobility of SMC1$^{4E}$ DNA, consistent with the hypothesis that cohesin$^{EQ/EQ}$ and cohesin-SMC1$^{4E}$ generated negatively and positively supercoiled plasmids in the presence of topoisomerase I, respectively. Representative image from 2 independent experiments. **(E)** Negatively supercoiled plasmid DNA was relaxed by incubating it with increasing concentrations of *E. coli* TopA (0, 0.28, 0.83, 2.5, 5 units). Representative image from 2 independent experiments.

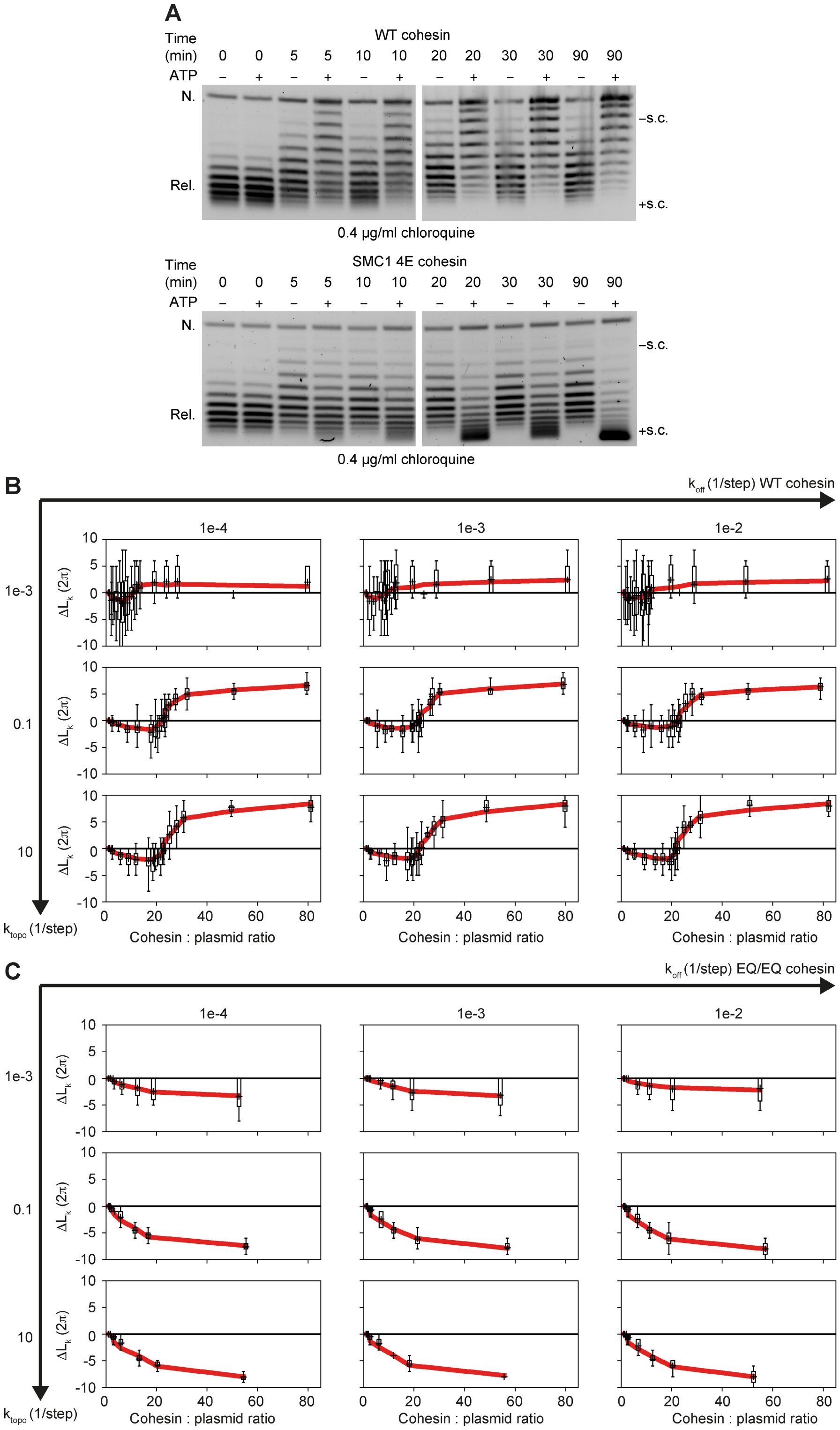


**Fig. S5. Loop extrusion-induced torque simulations can recapitulate the transition from negative to positive supercoiling chirality at high cohesin:plasmid DNA ratios. (A)** Human topoisomerase I DNA supercoiling assay in the presence or absence of the indicated proteins and ATP. Cohesin was added at a 15:1 molar ratio relative to plasmid DNA. NIPBL-MAU2 was added to all reactions at a 30:1 molar ratio relative to plasmid DNA. Reactions were terminated at the times indicated. Representative images from 3 independent experiments. **(B)** Simulations of wild type cohesin as in Fig. 4B for varying cohesin off-rates $k_{off}$ and topoisomerase rates $k_{topo}$ over two and five orders of magnitude, respectively. Bar plots represent the mean value (+ sign), whiskers denote the span of the data. The red line is a guide to the eye and was constructed by applying a Savitzky-Golay filter of window length 5 and order 1. Data are from 20 independent simulations. **(C)** As in (B) except for cohesin$^{EQ/EQ}$.

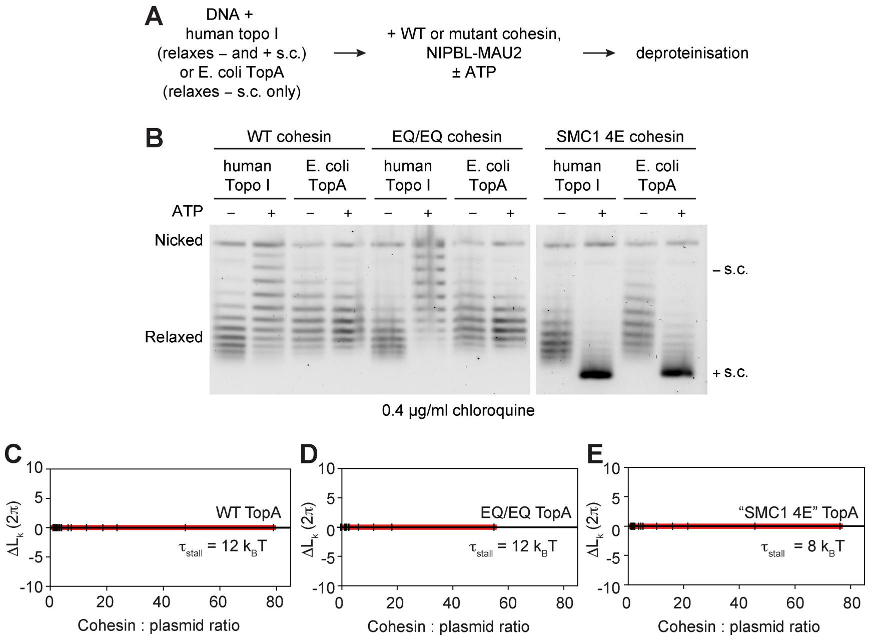


**Fig. S6. Wild type cohesin and cohesin$^{EQ/EQ}$ protect supercoils, but cohesin-SMC1$^{4E}$ does not. (A)** Experiment outline. **(B)** DNA supercoiling assay in the presence or absence of the indicated proteins and ATP. Cohesin was added at a 15:1 molar ratio relative to plasmid DNA. NIPBL-MAU2 was added to all reactions at a 30:1 molar ratio relative to plasmid DNA. Representative images from 2 independent experiments. **(C)** Simulation of the number and chirality of linking numbers removed (ΔLk) by *E. coli* TopA at the indicated wild type cohesin:plasmid ratios. *E.coli* TopA was simulated instead of human topo I by allowing that TopA can only resolve negative supercoils when the supercoiling density is s < 0.065. Bar plots represent the mean value (+ sign), whiskers denote the span of the data. The red line is a guide to the eye and was constructed by applying a Savitzky-Golay filter of window length 5 and order 1. Data are from 50 independent simulations. **(D)** As (C) but cohesin$^{EQ/EQ}$ was simulated by allowing only a single step per cohesin. **(E)** As in (C) except cohesin-SMC1$^{4E}$ was simulated by setting $t_{stall}$ at 8 $k_BT$. Note that the simulated outcome (no supercoiling) does not match the experimental outcome (positive supercoiling).

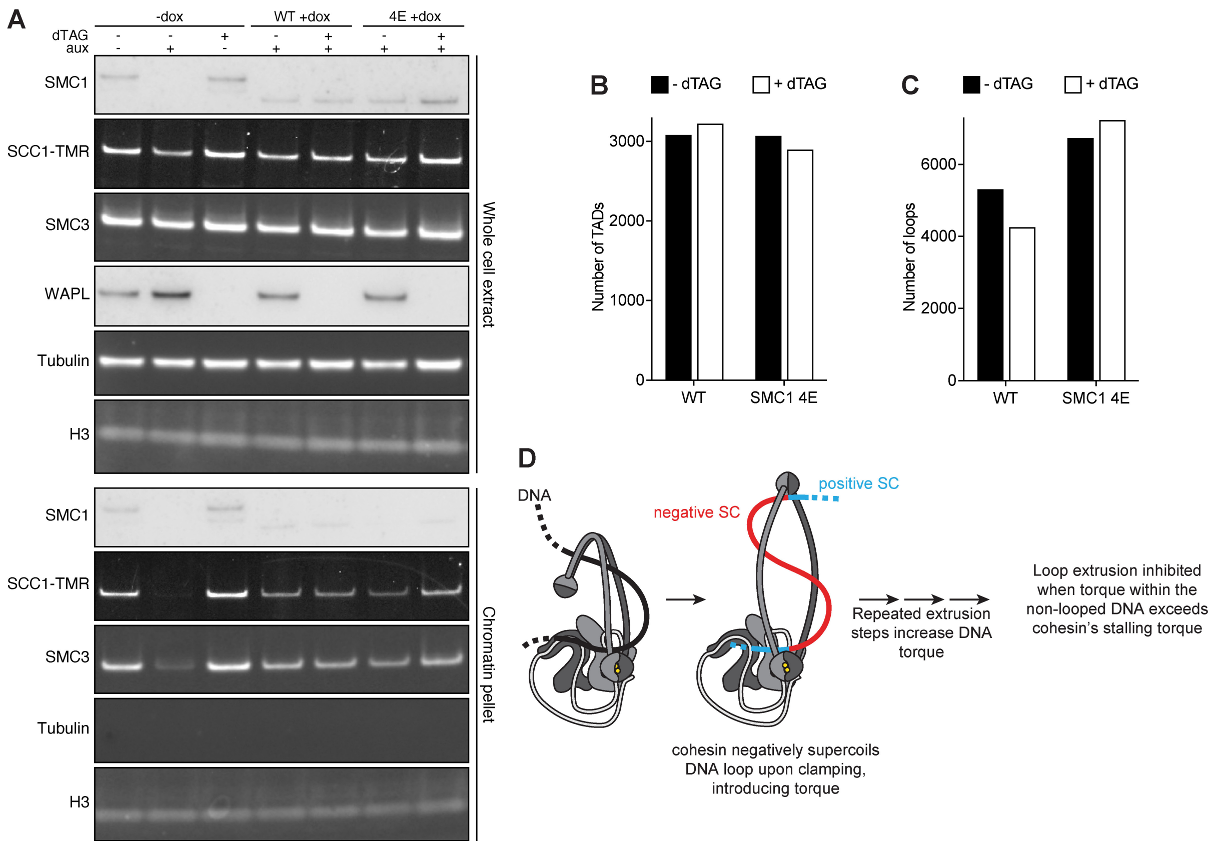


**Fig. S7. Characterization of HeLa cell lines expressing wild type SMC1 and SMC1$^{4E}$. (A)** Immunoblot analysis of whole-cell and chromatin extracts isolated from cells optionally treated with auxin, dTAG and doxycycline to degrade endogenous SMC1, degrade endogenous WAPL, and induce expression of wild type SMC1 or SMC1$^{4E}$, respectively. **(B and C)** Quantification of number of (B) TADs and (C) loops. **(D)** Model. Cohesin forms a negatively supercoiled DNA loop upon clamping, which introduces torque into both the loop and the non-looped DNA segments. Loop extrusion stalls once torque within the non-looped DNA exceeds a certain value. The torque required to stall cohesin is reduced in the presence of DNA clamp mutations.

| Assumption | Description |
|---|---|
| 1 | Cohesin binds to DNA with an on-rate of $1x10^{-5}$ to $1x10^{2}$ $s^{-1}$. |
| 2 | DNA-bound cohesin extrudes one step roughly every 0.5 s. |
| 3 | Cohesin binds uniformly along plasmid DNA in the absence of supercoiling. If supercoiling has accumulated, cohesin binds to a positively supercoiled segment with a 79 % probability(*31*). |
| 4 | Cohesin dissociates from DNA with a rate $k_{off.}$ |
| 5 | Cohesin moves directionally and can switch directions with a rate constant of $k_{switch}$=0.025 $s^{-1}$ (i.e., roughly once or twice per minute). Direction lifetimes are drawn from an exponential distribution with mean $1/k_{switch}$. |
| 6 | Each loop extrusion step enlarges the DNA loop by 150 bp (*50*). |
| 7 | Each loop extrusion step adds a negative twist of -0.6 into the DNA loop and a compensatory +0.6 twist or writhe into the non-looped DNA segment (Janissen et al., 2024). |
| 8 | Supercoiling transferred from the non-looped DNA segment into the loop in each step occurs according to the relative supercoiling per length in the non-looped segment and the loop extrusion step size. |
| 9 | Loop extrusion stalls when the absolute value of the torque within the looped or non-looped DNA segment reaches a stalling torque $\tau_{stall}$ = 12 $k_BT$. |
| 10 | Human topoisomerase I binds with uniform probability across the plasmid and removes both positive and negative supercoils in the respective DNA segment at the same rate $k_{topo.}$ |

**Table S1.**
Protocol and parameters used in the plasmid supercoiling simulations.

| No. | Name | Internal ref. | Vector | Ref. |
|---|---|---|---|---|
| 1 | SMC1, SMC3-FLAG | c190 (FW) | pBig1a | (*10*) |
| 2 | $SCC1^{R172A/D279A/R450A}$-HALO, HIS-STAG1 | c350 | pBig1b | this study |
| 3 | SMC1, SMC3-FLAG, $SCC1^{R172A/D279A/R450A}$-HALO, HIS-STAG1 | c357 | pBig2ab | this study |
| 4 | FLAG-HALO-NIPBL-HIS, MAU2 | c245/LC50A | pLib | (*10*) |
| 5 | PDS5A-HALO-HIS | c151 | pLib | (*10*) |
| 6 | $SMC1^{K38A}$, $SMC3^{K38A}$-FLAG | c130 | pBig1a | (*10*) |
| 7 | $SMC1^{E1157Q}$, $SMC3^{E1144Q}$-FLAG | c131 | pBig1a | (*10*) |
| 8 | $SMC1^{S1129R}$, $SMC3^{S1116R}$-FLAG | c367 | pBig1a | this study |
| 9 | $SMC1^{4E(K52E/R57E/K59E/R62E)}$, SMC3-FLAG | BB21/238step1 | pBig1b | (*12*) |
| 10 | $SMC1^{\Delta 58\text{-}62}$, SMC3-FLAG | BB20/262step1 | pBig1b | (*12*) |
| 11 | SMC1, $SMC3^{4E(R57E/R61E/K105E/K106E)}$-FLAG | BB23/239step1 | pBig1b | (*12*) |
| 12 | SMC1, $SMC3^{2E(R57E/R61E)}$-FLAG | BB22/285step1 | pBig1b | (*12*) |
| 13 | $SMC1^{hinge3A(K551A/R554A/K561A)}$, $SMC3^{hinge1A(R644A)}$-FLAG | BB24/251step1 | pBig1b | (*12*) |
| 14 | $SMC1^{K52E}$, SMC3-FLAG | c374 | pBig1a | this study |
| 15 | $SMC1^{R57E}$, SMC3-FLAG | c375 | pBig1a | this study |
| 16 | $SMC1^{K59E}$, SMC3-FLAG | c376 | pBig1a | this study |
| 17 | $SMC1^{R62E}$, SMC3-FLAG | c377 | pBig1a | this study |
| 18 | HIS-YBBR-FLAG-TOPOISOMERASEI | c234 | pLib | this study |
| 19 | SMC1, SMC3-FLAG, HIS-SCC1(TEV)-HALO | s15c | pBig2ab | this study |
| 20 | HIS-STAG1 | c179 (FW) | pLib | (*10*) |
| 21 | $SMC1^{4E\text{-}EQ(K52E/R57E/K59E/R62E/E1157Q)}$, $SMC3^{E1144Q}$-FLAG | c366 | pBig1a | this study |
| 22 | $SMC1^{4E\text{-}SR(K52E/R57E/K59E/R62E/S1129R)}$, $SMC3^{S1116R}$-FLAG | c368 | pBig1a | this study |

**Table S2.**
Protein expression constructs used in this study.

**Movie S1. High-speed atomic force microscopy recording of purified plasmid DNA following incubation with topoisomerase I and ATP.**

**Movie S2. High-speed atomic force microscopy recording of purified plasmid DNA following incubation with topoisomerase I, wild type cohesin, NIPBL-MAU2 and ATP.**

**Movie S3. Example simulation of plasmid supercoiling by cohesin.** During this simulation, 14 cohesin complexes extruded the plasmid. Description of graphs from top left to bottom right: (i) Loop positions on a plasmid of 4000 bp length (using periodic boundary conditions). The non-extruded part of the plasmid is colored light blue in all panels and is labeled 'loop 0' with loop anchors at 0 bp and 4000 bp. Z-loops are drawn uniformly in grey. (ii) The net topoisomerase-removed ΔLk as a function of loop extrusion (LE) steps. (iii) The torque of individual segments of the plasmid (backbone, loops, nested loops, and Z-loops) as a function of LE steps. (iv) The black line refers to the left y-axis and shows the fraction of the plasmid with positive twist (Tw > 0). The red and blue lines refer to the right y-axis and display the fraction of the plasmid with Tw > 1 and Tw < -1, respectively, i.e. the positively and negatively supercoiled fraction of the plasmid which can be relaxed by topoisomerase. (v) The momentary length of loops and Z-loops is shown on the y-axis and the x-axis is the coordinate along the plasmid with periodic boundary conditions. The loop anchor positions are the same as in panel (i). Note, however, that the loop length can be different from the difference between loop anchors if cohesin complexes extrude nested loops or Z-loops. (vi) The momentary twist for each segment displayed as in (v). (vii) The torque of each segment displayed as in (v). (viii) The maximum ΔLk that topoisomerase can remove in every loop segment. This panel displays the nearest smaller integer of the values displayed in panel (vi). The first part of the video is frequently paused and annotations were added to describe elements of the simulation as well as the progression and interaction of cohesin with topoisomerases. The simulation then proceeds and its result is related to the graph shown in Figure 4B (14 cohesin complexes extruded the plasmid in this example). Following this, the simulation is repeated without annotation.

**Movie S4. Additional example simulation of plasmid supercoiling by cohesin.** During this simulation, 43 cohesin complexes extruded the plasmid. For a description of graphs, see the legend to Movie S3.
